# Efficacy and acceptability of non-invasive brain stimulation for the treatment of adult unipolar and bipolar depression: A systematic review and meta-analysis of randomised sham-controlled trials

**DOI:** 10.1101/287656

**Authors:** Julian Mutz, Daniel R. Edgcumbe, Andre R. Brunoni, Cynthia H.Y. Fu

**Author notes:** Author for Correspondence: Julian Mutz, Department of Epidemiology and Biostatistics, School of Public Health, Faculty of Medicine, Imperial College London, United Kingdom; Norfolk Place, London W2 1PG, United Kingdom.

## Abstract

We examined the efficacy and acceptability of non-invasive brain stimulation in adult unipolar and bipolar depression. Randomised sham-controlled trials of transcranial direct current stimulation (tDCS), transcranial magnetic stimulation (TMS) and theta-burst stimulation (TBS), without co-initiation of another treatment, were included. We analysed effects on response, remission, all-cause discontinuation rates and continuous depression severity measures. Fifty-six studies met our criteria for inclusion (*N* = 3,058, mean age = 44.96 years, 61.73% female). Response rates demonstrated efficacy of high-frequency rTMS over the left DLPFC (OR = 3.75, 95% CI [2.44; 5.75]), right-sided low-frequency rTMS (OR = 7.44, 95%CI [2.06; 26.83]) bilateral rTMS (OR = 3.68,95%CI [1.66; 8.13]), deep TMS (OR = 1.69, 95%CI [1.003; 2.85]), intermittent TBS (OR = 4.70, 95%CI [1.14; 19.38]) and tDCS (OR = 4.17, 95% CI [2.25; 7.74]); but not for continuous TBS, bilateral TBS or synchronised TMS. There were no differences in all-cause discontinuation rates. The strongest evidence was for high-frequency rTMS over the left DLPFC. Intermittent TBS provides an advance in terms of reduced treatment duration. tDCS is a potential treatment for non-treatment resistant depression. To date, there is not sufficient published data available to draw firm conclusions about the efficacy and acceptability of TBS and sTMS.

**Highlights:**

- Response, remission, all-cause discontinuation rates and continuous post-treatment depression scores were examined
- Several non-invasive brain stimulation treatments seem efficacious across different outcome metrics
- All-cause discontinuation rates indicate no differences between sham and active treatment

## Introduction

Major depression is prevalent^1^ and associated with considerable disease burden^2^. Its course is often recurrent and may become chronic with relapse rates within one year of remission ranging from 35% to 80%^3,4^. The most common treatments are pharmacological and psychological therapies. Yet, even with a full course of treatment, at least one third of patients fail to achieve remission^5^. Non-invasive neurostimulation therapies, such as transcranial magnetic stimulation (TMS) and transcranial electrical stimulation (tES), offer a potential alternative or add-on treatment strategy.

TMS was originally introduced as a tool for investigating and mapping cortical functions and connectivity^6^. TMS utilises intense, rapidly-changing electromagnetic fields generated by a coil of wire near the scalp and allows for a mostly undistorted induction of an electrical current to alter neural activity in relatively focal, superficial areas of the brain. Standard TMS involves single or paired pulses, while repetitive transcranial magnetic stimulation (rTMS) involves the delivery of repeated pulses which enable the prolonged modulation of neural activity. Depending on the stimulation frequency, rTMS can increase or decrease cortical excitability. The prevailing hypothesis is that the aftereffects of high-frequency (usually 10Hz or higher) stimulation are excitatory while those of low-frequency (≤1Hz) stimulation are inhibitory^7^.

The rationale for using rTMS to treat depressive illness comes from clinical symptomatology and neuroanatomy as well as neuroimaging studies indicating functional impairments in prefrontal cortical and limbic regions^8^. In 2008, the US Food and Drug Administration (FDA) approved the first rTMS device for the treatment major depressive disorder (MDD) in which there was poor response to at least one pharmacological agent in the current episode^9^, and its clinical utilisation has increased since^10^.

As stimulation at high frequencies can be uncomfortable during the initial stimulation period, low-frequency rTMS may minimise the occurrence of undesired side effects, namely headaches and scalp discomfort, and may be associated with fewer adverse events, for instance by lowering the risk for developing seizures^11^.

Bilateral applications of rTMS have also been developed: simultaneous stimulation over the left and right DLPFC (rDLPFC) or stimulation over one side followed by stimulation of the other side. These applications were hypothesised to be potentially additive or synergistic to reinstate any imbalance in prefrontal neural activity^12^. Moreover, there may be a selective unilateral response and the likelihood for a clinical response may increase by providing both types of stimulation^13^.

Technical and methodological efforts to improve the antidepressant efficacy of TMS have led to several alternative treatment protocols. Deep TMS (dTMS) was FDA-approved in 2013, which is able to stimulate larger brain volumes and deeper structures^14^ that could be more directly relevant in the pathophysiology of depression (e.g., reward-mediating pathways and areas connected to the subgenual cingulate cortex)^8,15,16^.

Another recent modification is theta burst stimulation (TBS)^17^, which is a patterned form of TMS pulse delivery that utilises high and low frequencies in the same stimulus train. TBS delivers bursts of three at a high frequency (50Hz) with an inter-burst interval of 5Hz in the theta range at 5Hz. Two different protocols are utilised: continuous theta burst stimulation (cTBS), which delivers 300 or 600 pulses without interruption, and intermittent theta burst stimulation (iTBS), which delivers 30 pulses every 10 seconds for a duration of 190 seconds, totalling 600 pulses^18^. It is suggested that cTBS reduces cortical excitability while iTBS increases it, mimicking the processes of long-term potentiation and long-term depression, respectively^17^. Notably, there is some debate as to whether prolonged stimulation periods reverse the hypothesised effects of TBS^19^, while there is also support for a dose-response relationship for iTBS^20^.

The main advantages of TBS are its reduced administration time, which is typically less than five minutes as opposed to 20–45 minutes for conventional rTMS, and the lower intensity needed to produce lasting neurophysiological effects as TBS is typically administered at 80% of the resting motor threshold (rMT) and might be more comfortable than stimulation at higher intensities typically used with standard rTMS.

Synchronised TMS refers to magnetic low-field synchronised stimulation (sTMS), a new treatment paradigm that involves rotating spherical rare-earth (neodymium) magnets positioned sagittally along the midline of the scalp, which deliver stimulation synchronised to an individual’s alpha frequency^21^. The magnets are positioned to provide a global magnetic field distributed broadly across the midline cortical surface (one magnet over the frontal polar region, one magnet over the top of the head, and one magnet over the parietal region). The rationale for sTMS synchronised to an individual’s alpha frequency is the observation that one mechanism of action of rTMS is the entrainment of oscillatory activity to the programmed frequency of stimulation, thereby resetting thalamo-cortical oscillators and restoring normal endogenous oscillatory activity^22^. This modification of TMS may be associated with fewer treatment-emergent adverse and side effects because it does not cause neural depolarisation. It also uses less energy than conventional rTMS as it utilises sinusoidal instead of pulsed magnetic fields, which require less than 1% of the energy needed for conventional rTMS and may thus be less expensive.

Access and costs are among the major impediments to a more widespread use of rTMS, although costs may be lower for TBS and sTMS. A less expensive technique is transcranial electrical stimulation (tES). Its most commonly used protocol, transcranial direct current stimulation (tDCS), was reappraised as a tool in research through the work of Priori et al.^23^ and Nitsche and Paulus^24^. tDCS involves the application of a low-amplitude electrical direct current through surface scalp electrodes to superficial areas of the brain. While it does not directly trigger action potentials, it modulates cortical excitability by shifting the neural membrane resting potential and these effects can outlast the electrical stimulation period^25^. The direction of such excitability changes may depend on the polarity of the stimulation: anodal stimulation is hypothesised to cause depolarisation and an increase in neural excitability, whereas cathodal stimulation causes hyperpolarisation and a decrease in cortical excitability^26,27^.

The advantages of tDCS compared to TMS include its ease of administration, being much less expensive, its more benign side effect profile, and its portability which could potentially be used in the home environment^28^.

We sought to perform a systematic review and meta-analysis of the antidepressant efficacy and acceptability of non-invasive neuromodulation in treating a current depressive episode in unipolar and bipolar depression from randomised sham-controlled trials. The only study to date that evaluated the efficacy of a range of rTMS techniques is Brunoni et al.'s network meta-analysis^29^. However, the analysis had included trials that had co-initiated other treatments (e.g. sleep deprivation and TMS); trials which had not included a sham treatment; had not separated the TBS modifications; and had not included any age-related exclusion criteria. Also, tDCS trials were not included in that meta-analysis. We sought to address these limitations by including only trials with randomised allocation to active or sham treatments, excluding studies which had co-initiated another treatment, and limiting our sample to the adult age range as geriatric depression may impact on efficacy.

## Materials and Methods

### Search strategy and selection criteria

We followed the Preferred Reporting Items for Systematic Reviews and Meta-Analyses (PRISMA) guidelines^30^. A systematic search of the Embase, Medline, and PsycINFO databases was performed from the first date available to 1st May 2018 (Figure 1). The following search terms were used: (bipolar disorder OR bipolar depression OR major depression OR unipolar depression OR unipolar disorder) AND (transcranial direct current stimulation OR tDCS OR transcranial magnetic stimulation OR TMS OR theta burst stimulation OR TBS OR sTMS OR dTMS), limiting searches to studies in humans and English-language publications. Reference lists of included papers and of recent systematic reviews and meta-analyses (Supplementary Material 1) were screened for further studies. This study has not been previously registered.

**Figure 1.**
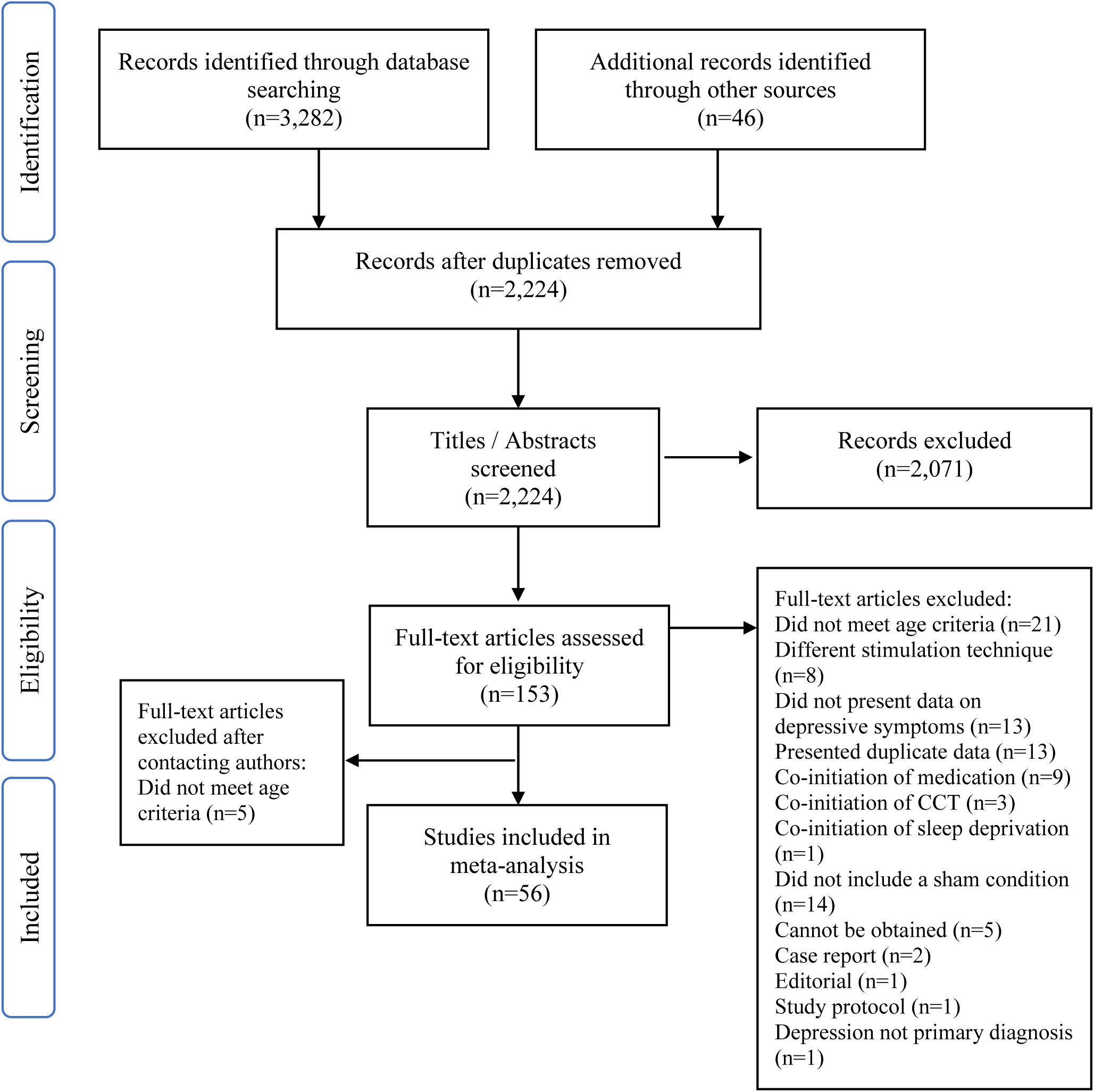
Caption: Preferred Reporting Items for Systematic Reviews and Meta-Analyses (PRISMA) flow diagram of literature search.

Inclusion criteria were: 1) adults aged 18 - 70 years; 2) DSM or ICD diagnosis of MDD or bipolar disorder currently in a major depressive episode; 3) randomised sham-controlled trials, which utilised a parallel-group or cross-over design; 4) clinician-administered depression rating scale, Hamilton Depression Rating Scale (HDRS)^31^ or Montgomery-Åsberg Depression Rating Scale (MADRS)^32^.

Exclusion criteria were: 1) primary diagnoses other than MDD or bipolar depression; 2) studies limited to a specific subtype of depression (e.g., postpartum depression or vascular depression) or in which a major depressive episode was a secondary diagnosis (e.g., fibromyalgia and major depression); 3) co-initiation of any other form of treatment, such as pharmacotherapy or cognitive control training.

### Data analysis

The following sample characteristics were extracted: sex, age, hospitalisation status, whether patients with psychotic symptoms were excluded from the study, diagnosis, treatment strategy, and treatment resistance.

The following treatment-related parameters were extracted. For TMS: type of coil and sham procedure, coil location, stimulation frequency (Hz) for each site, stimulation intensity (percentage of the rMT), total number of pulses delivered, and number of treatment sessions. For TBS: data on the treatment protocol (iTBS, cTBS or bilateral TBS) were also recorded. For tDCS: location of the anode and cathode, electrode size (cm^2^), current intensity (mA) and density (mA/cm^2^), session duration, number of sessions, and duration of active stimulation in the sham condition.

The primary outcome measure was clinical response, defined as a ≥ 50% reduction in symptom scores at the primary study endpoint. Remission rates were the secondary outcome measure based on the definition provided by each study. If response or remission rates were reported for both HDRS and MADRS, data for the HDRS were selected to facilitate comparability between trials. If data for multiple versions of the HDRS were reported, the original 17-item version was selected. We also extracted baseline and post-treatment depression severity scores; the latter constituted our tertiary outcome measure. If available, the intention-to-treat (ITT) or modified intention-to-treat (mITT) data were preferred over data based only on completers. For cross-over trials, only data from the initial randomisation were used to avoid carry-over effects. Data presented in figures were extracted with WebPlotDigitizer (http://arohatgi.info/WebPlotDigitizer/app/). All-cause discontinuation rates were recorded separately for active and sham groups and were treated as a primary outcome measure of acceptability.

Data that could not be directly retrieved from the original publications were requested from the authors or searched for in previous systematic reviews and meta-analyses. For trials with more than two groups that could not be included as separate treatment comparisons, we combined groups to create single pair-wise comparisons.

For dichotomous outcome data, odds ratios (Mantel-Haenszel method) were used as an index of effect size. We also computed Hedge’s *g* to estimate the effect sizes for continuous post-treatment depression scores. A random-effects model was chosen as it was assumed that the underlying true effect size would vary between studies. A random-effects model provides wider confidence intervals than a fixed-effects model if there is significant heterogeneity among studies and thus tends to be more conservative in estimating summary effect sizes.

Contour-enhanced funnel plots^33^ were visually inspected to assess whether potential funnel asymmetry is likely to be due to statistical significance-based publication bias.

Heterogeneity between studies was assessed with the Qt statistic, which estimates whether the variance of effect sizes is greater than what would be expected due to sampling error. A *p* value smaller than .01 provides an indication for significant heterogeneity^34^. The I^2^ statistic was computed for each analysis to provide a descriptive measure of inconsistency across the results of individual trials included in our analyses. It provides an indication of what percentage of the observed variance in effect sizes reflects real differences in effect sizes as opposed to sampling error. Higgins et al.^35^ suggested that 25%, 50%, and 75% represent little, moderate, and high heterogeneity, respectively.

Where sufficient data were available, we conducted subgroup analyses to examine potential differences in antidepressant efficacy by clinical and study characteristics including diagnosis, whether the trial excluded patients with psychotic symptoms, hospitalization status and treatment resistance.

Analyses were conducted using the ‘meta’ package^36^ for RStudio (Version 0.98.932) and STATA (Version 13.1; StataCorp, 2013) was used for data processing.

The Cochrane tool for assessing risk of bias in randomised trials^37^ was used to evaluate included studies. Each trial received a score of low, high, or unclear risk of bias for each of the potential sources of bias. Two raters independently conducted the assessment of risk of bias.

## Results

### Overview

Fifty-six RCTs, consisting of 131 treatment arms met our criteria for inclusion (Figure 1, Supplementary Material 2). Overall, 66 treatment comparisons were included, total *N* = 3,058 patients (mean age = 44.96 years, 61.73% female) of whom *n* = 1,598 were randomised to active and *n* = 1,460 to sham treatments (Tables 1–4).

**Table 1.**
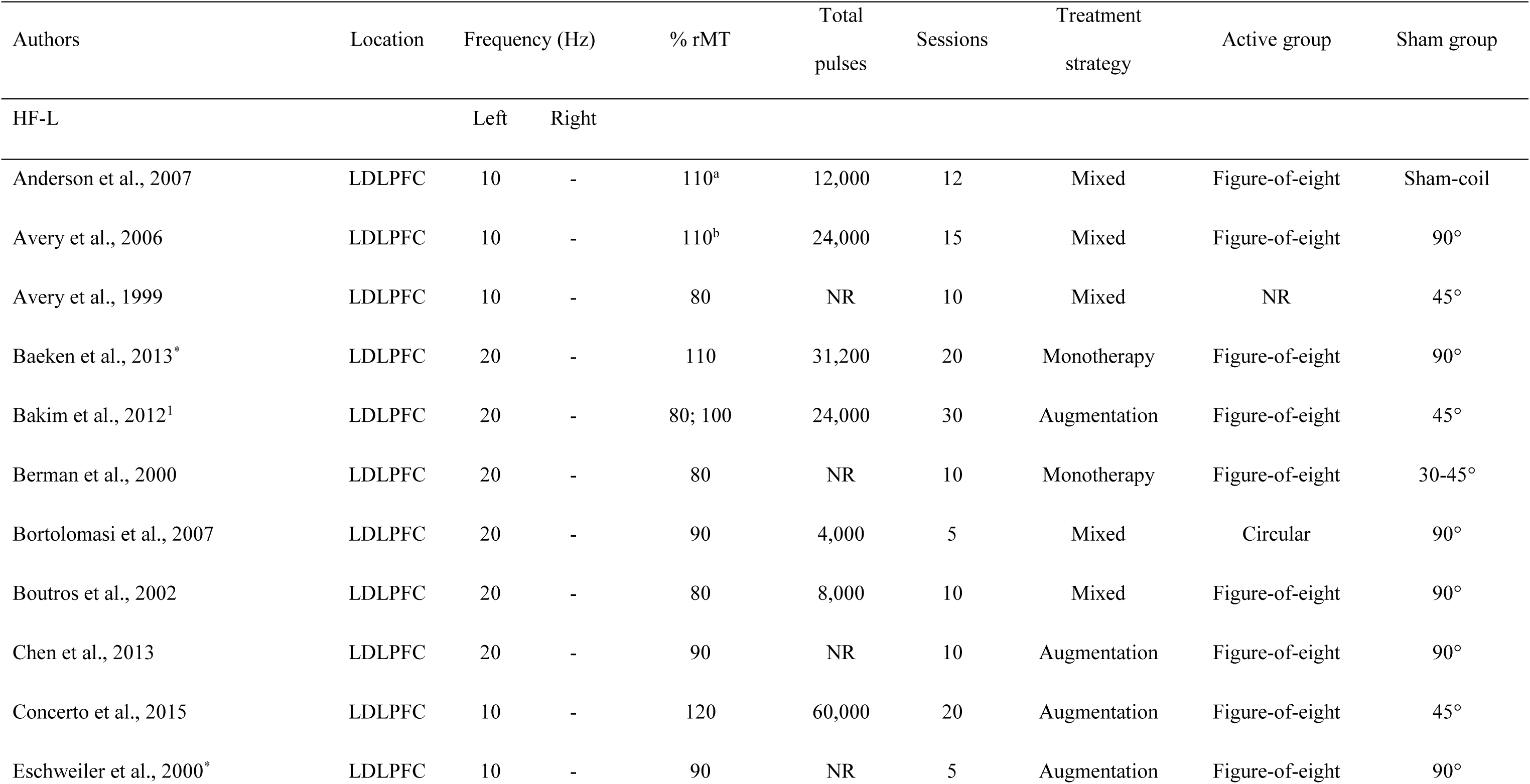
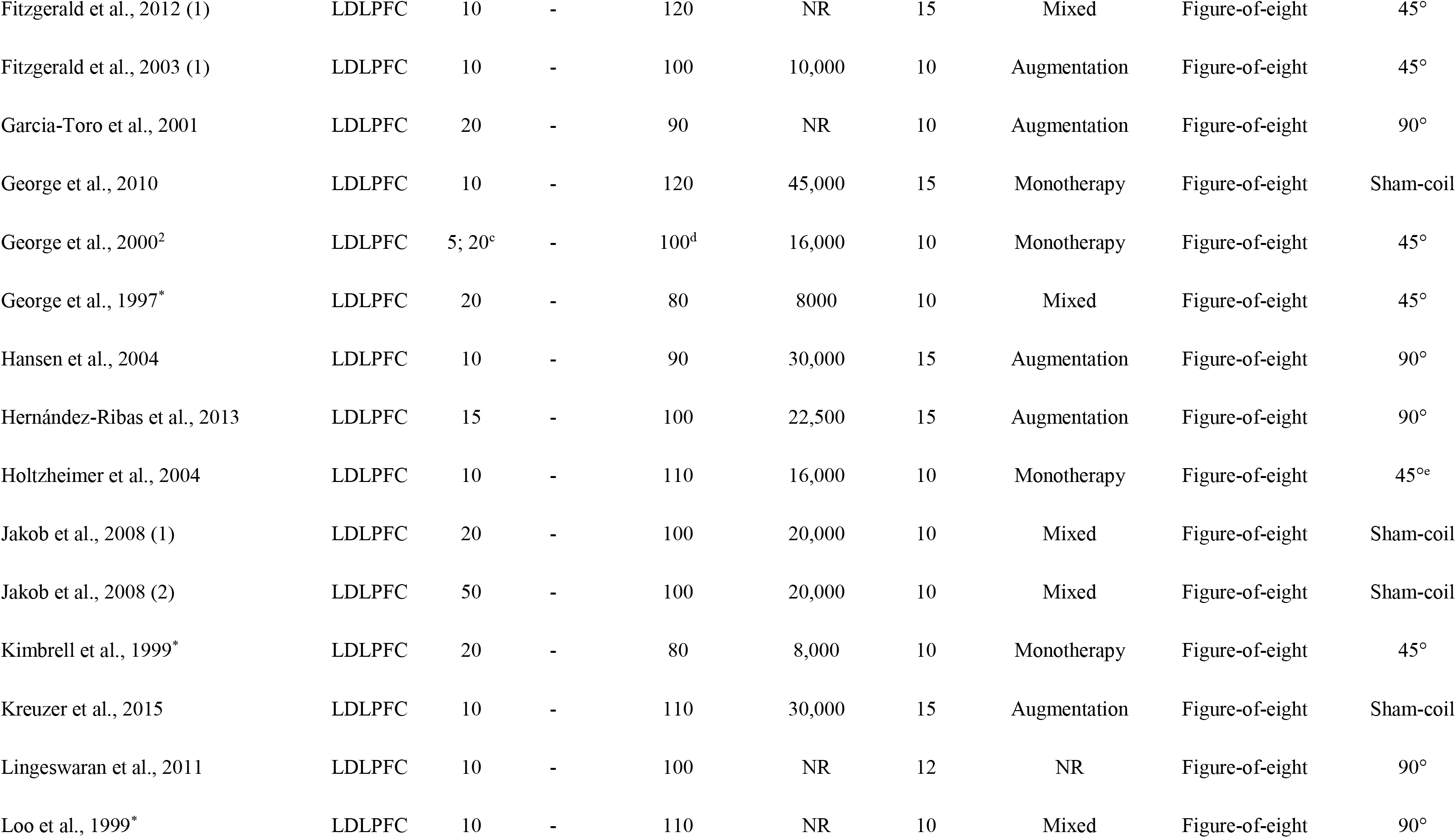
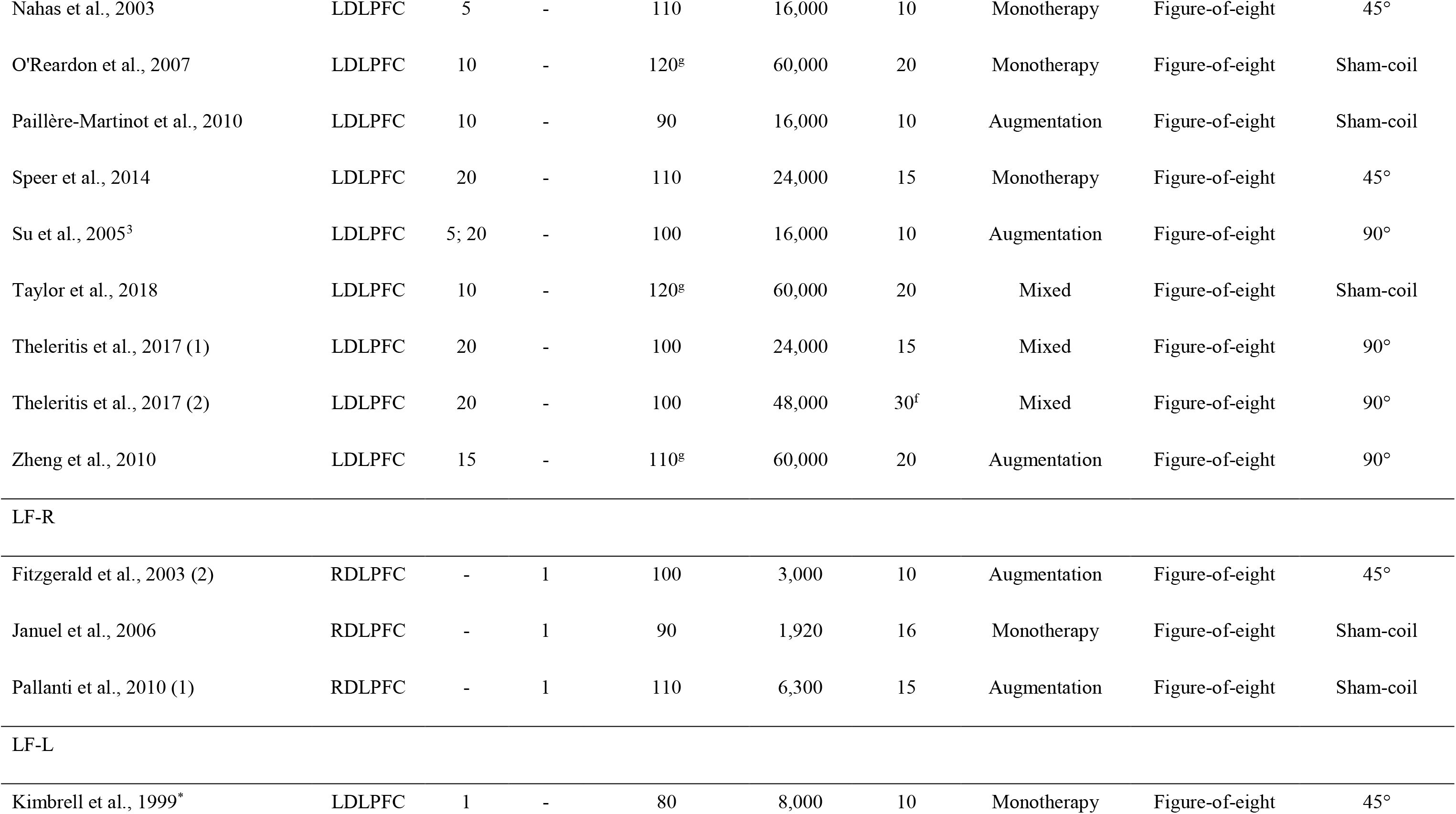
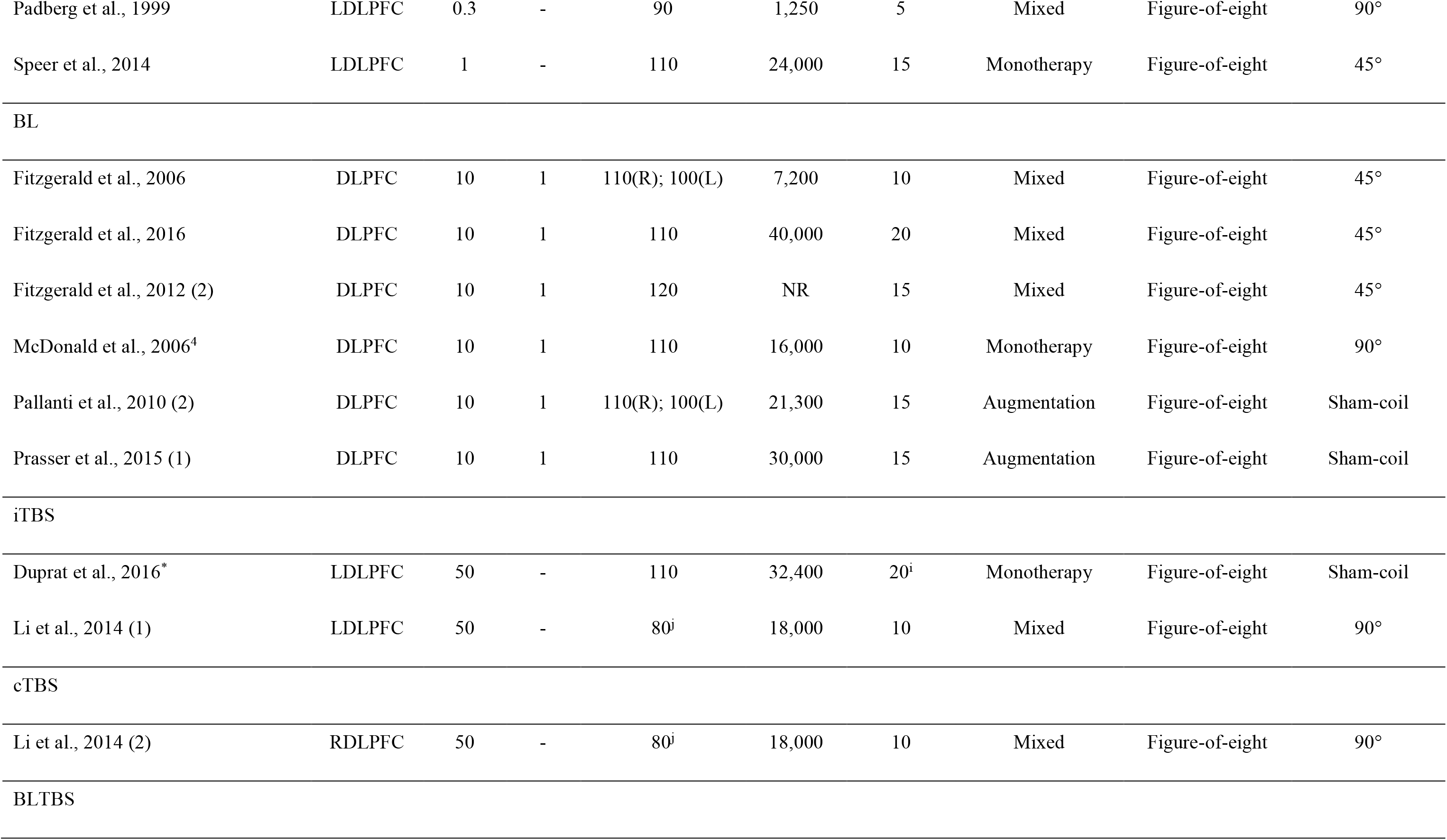
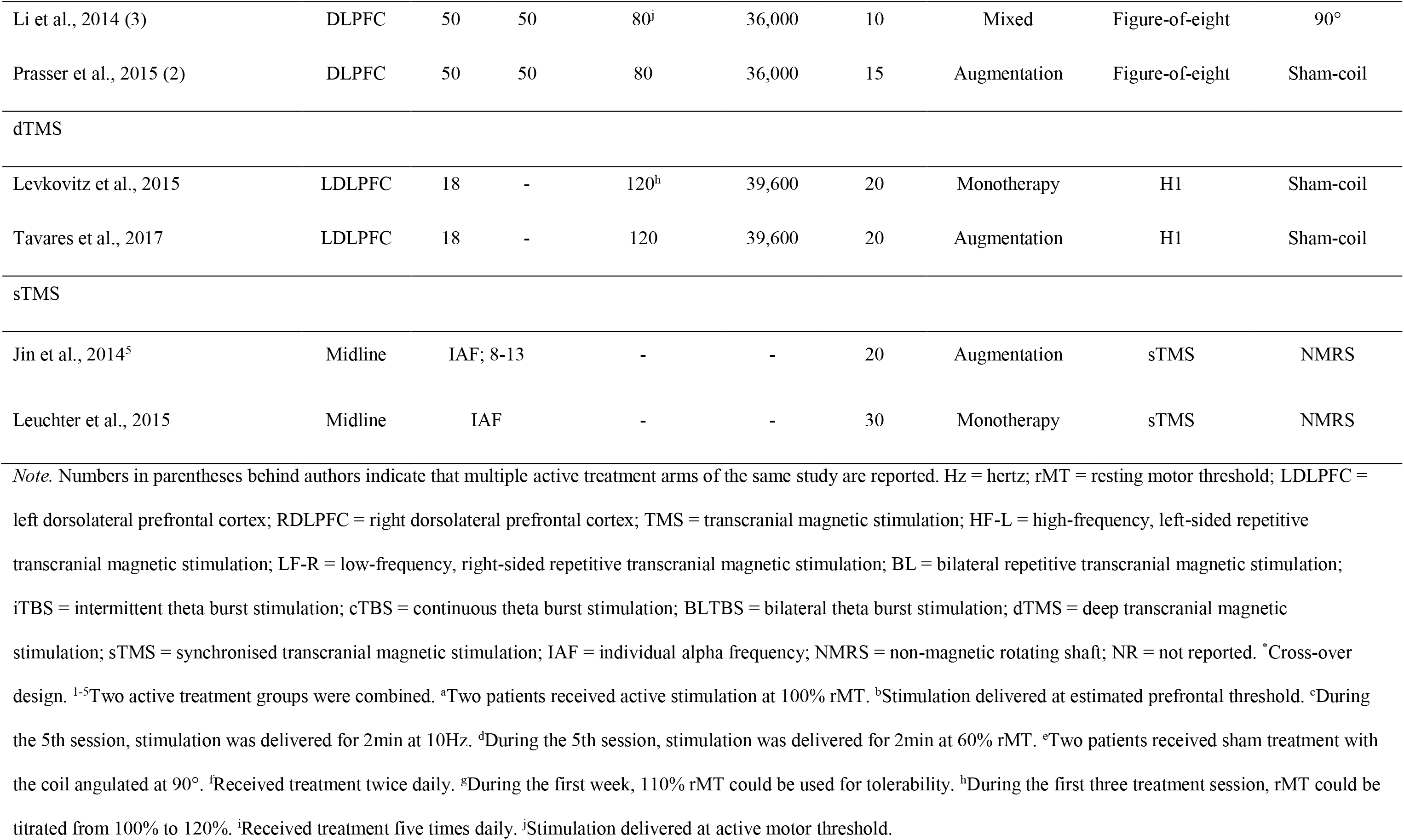
Treatment characteristics: TMS studies

**Table 2.**
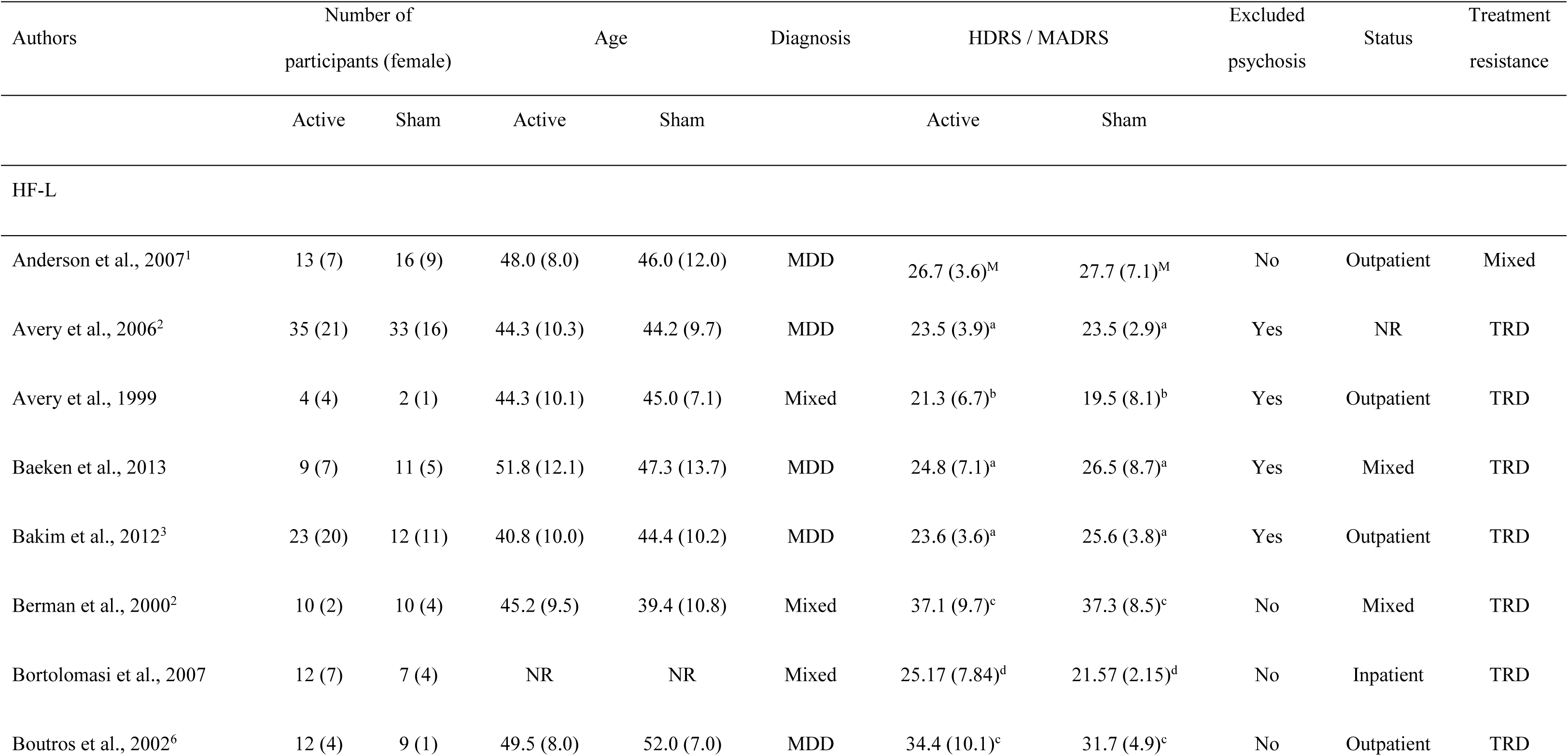
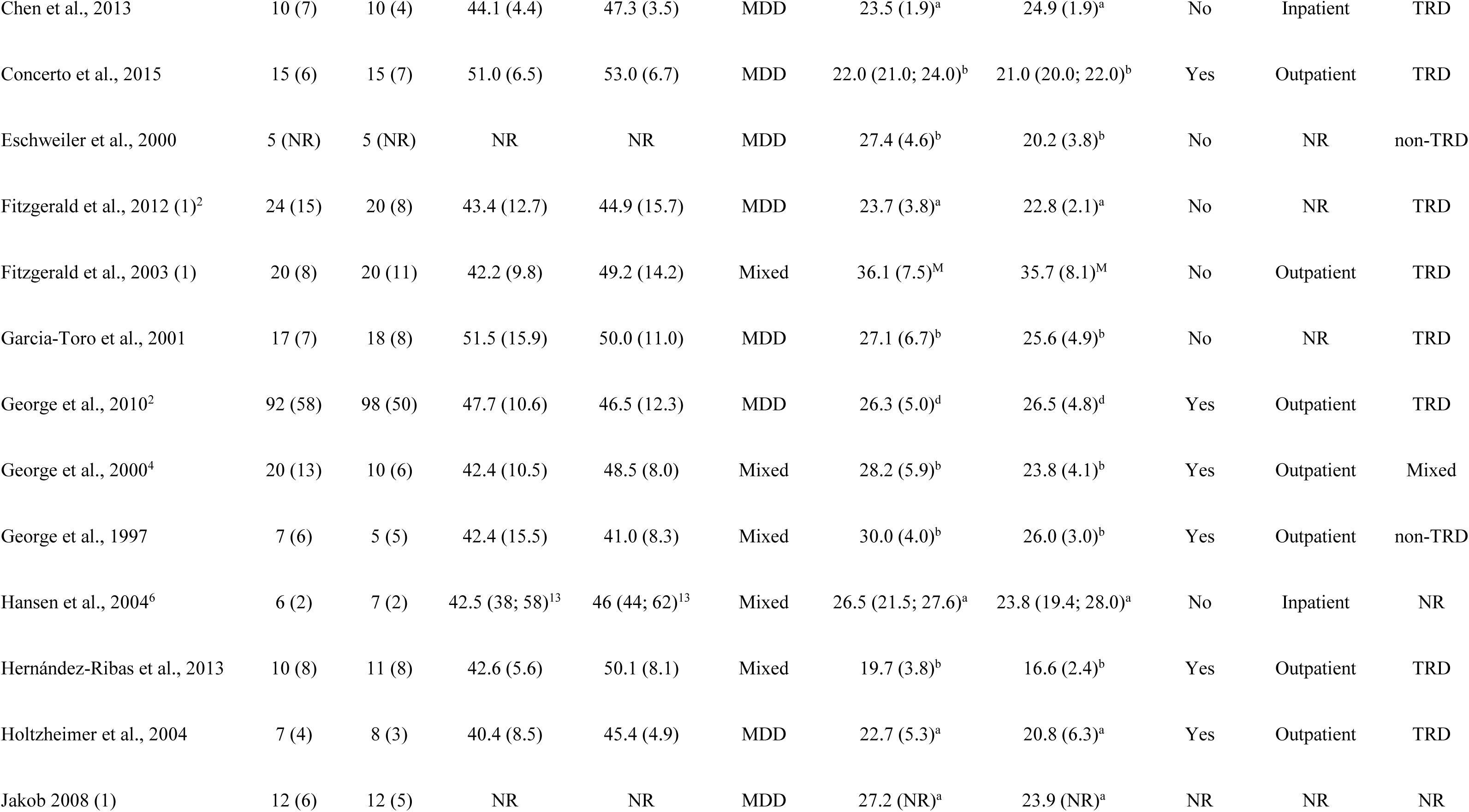
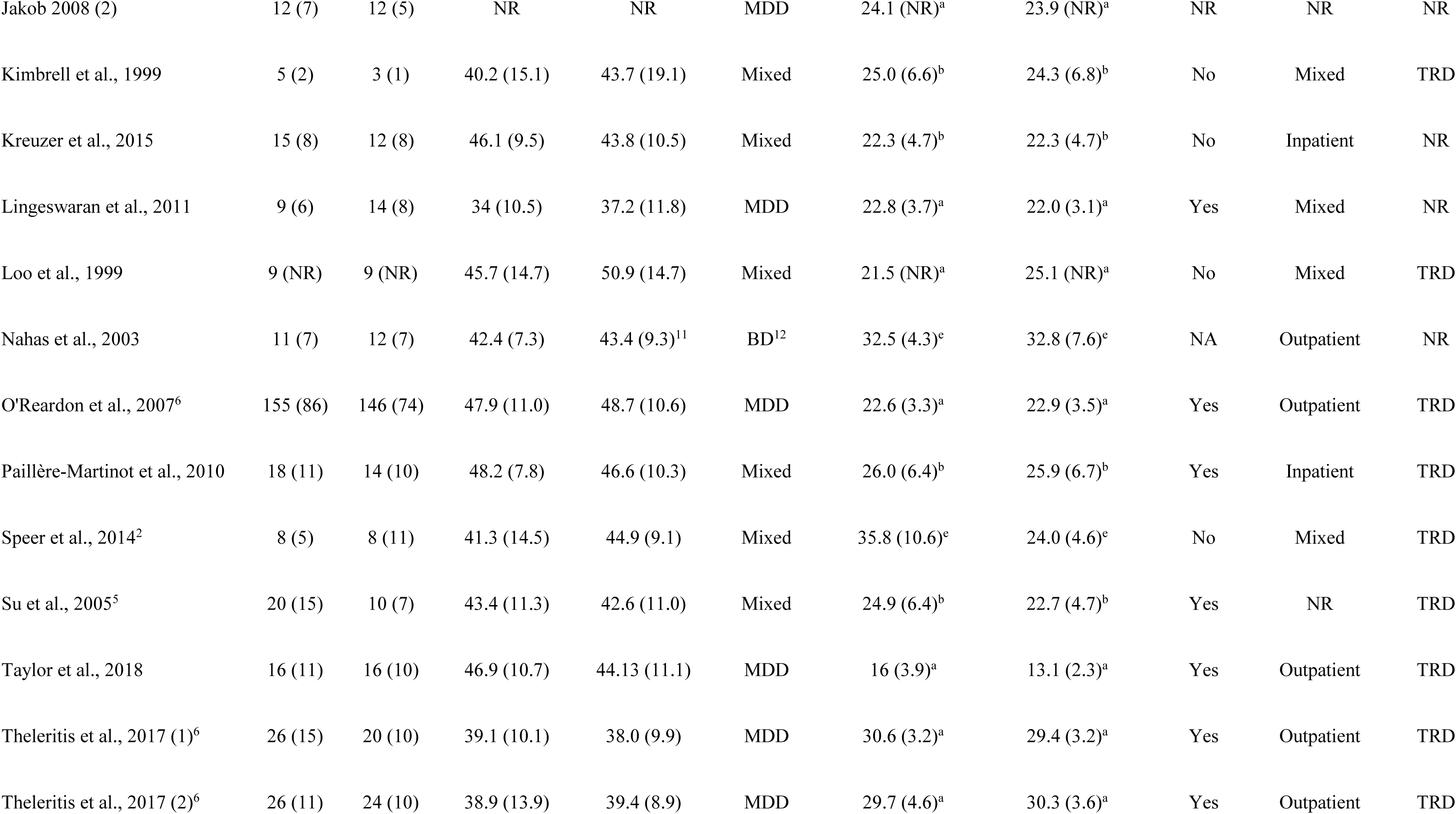
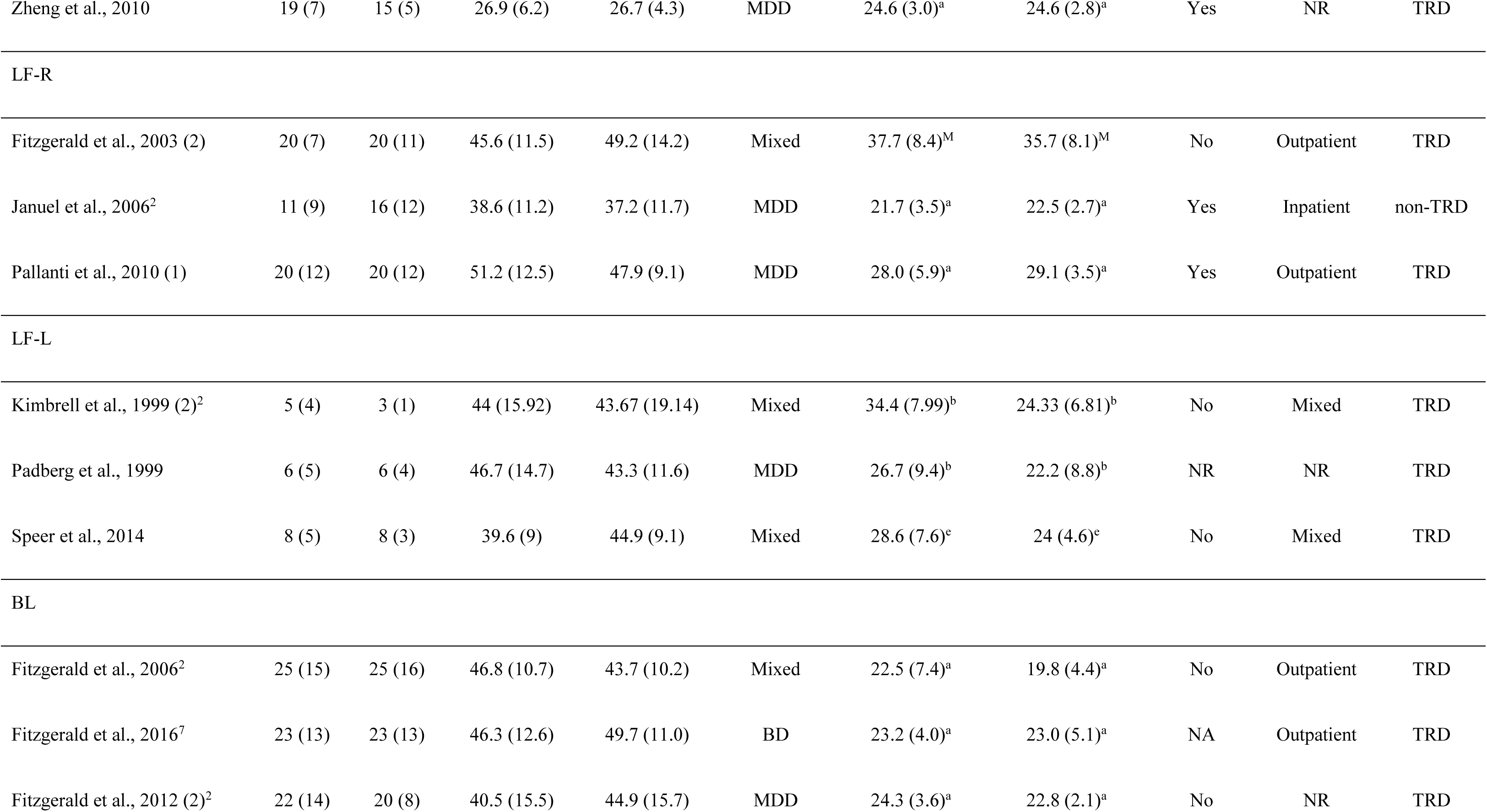
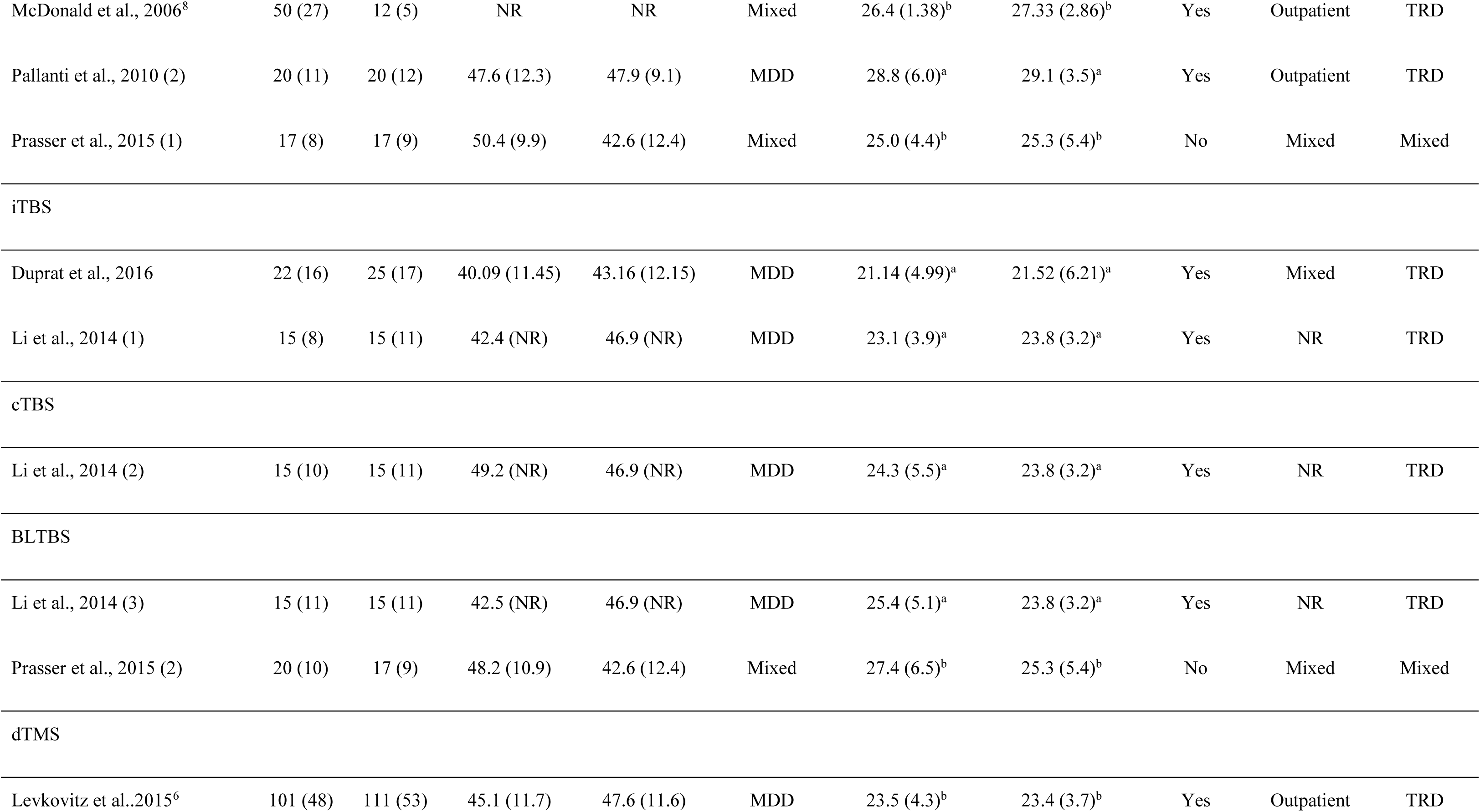
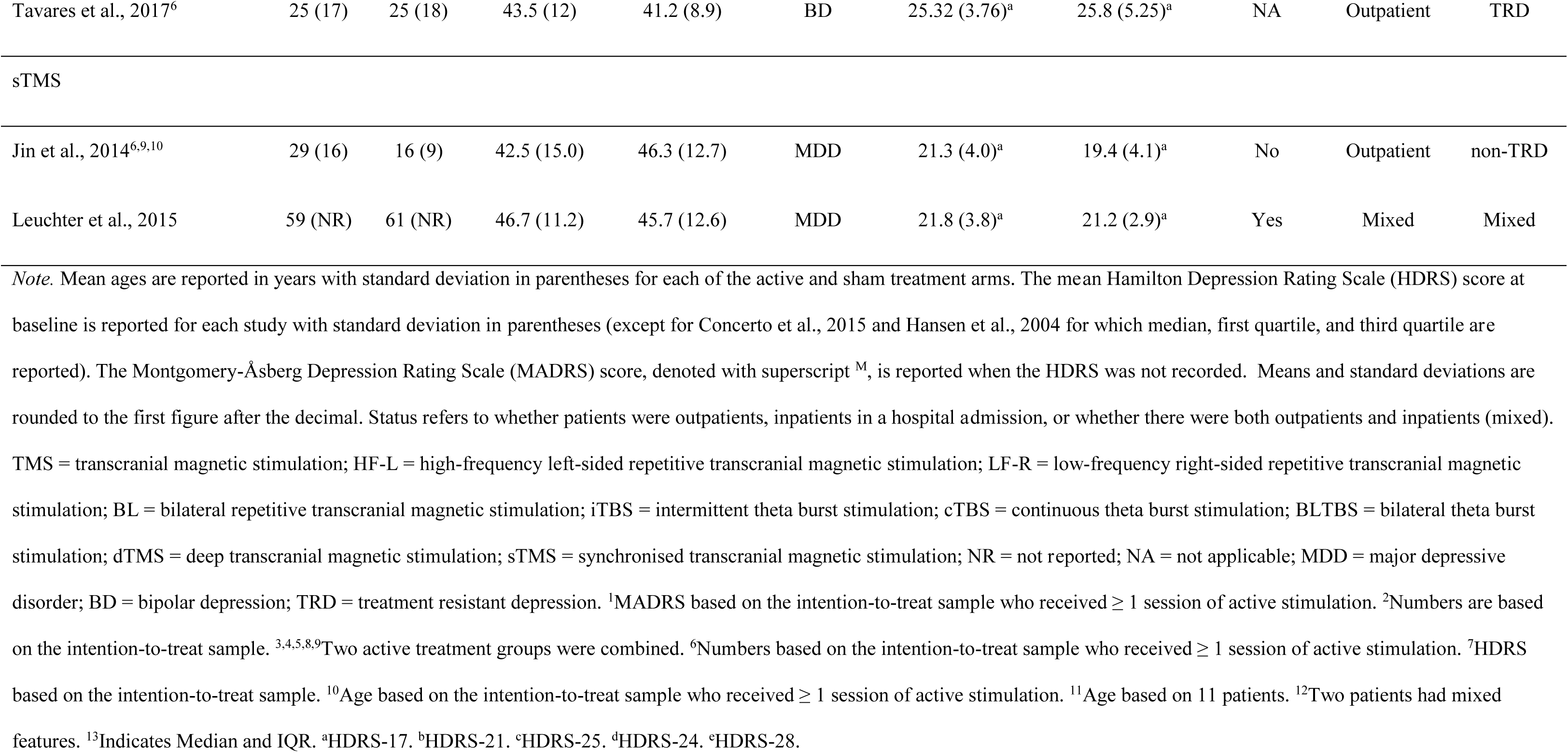
Sample characteristics: TMS studies

**Table 3.** Treatment characteristics: tDCS studies

| Authors | Location |  | Electrode size | Current strength | Current density | Session duration | Number of sessions | Treatment strategy | Sham stimulation |
| --- | --- | --- | --- | --- | --- | --- | --- | --- | --- |
|  | Anode | Cathode/Reference |  |  |  |  |  |  |  |
| Fregni et al., 2006a | F3 | FP2 | 35cm<sup>2</sup> | 1mA | 0.028 | 20min | 5 | Monotherapy | 05sec |
| Fregni et al., 2006b | F3 | FP2 | 35cm<sup>2</sup> | 1mA | 0.028 | 20min | 5 | Monotherapy | 05sec |
| Boggio et al., 2008<sup>1</sup> | F3 | FP2; Midline | 35cm<sup>2</sup> | 2mA | 0.057 | 20min | 10 | Monotherapy | 30sec |
| Loo et al., 2010 | pF3 | F8 | 35cm<sup>2</sup> | 1mA | 0.028 | 20min | 5 | Mixed | 30sec |
| Blumberger et al., 2012 | F3 | F4 | 35cm<sup>2</sup> | 2mA | 0.057 | 20min | 15 | Mixed | 30sec |
| Brunoni et al., 2013<sup>2</sup> | F3 | F4 | 25cm<sup>2</sup> | 2mA | 0.080 | 30min | 12 | Monotherapy | 60sec |
| Salehinejad et al., 2015 | F3 | F4 | 35cm<sup>2</sup> | 2mA | 0.057 | 20min | 22 | Monotherapy | 30sec |
| Salehinejad et al., 2017 | F3 | F4 | 35cm<sup>2</sup> | 2mA | 0.057 | 30min | 10 | Monotherapy | 30sec |
| Brunoni et al., 2017<sup>2</sup> | F3 | F4 | 25cm<sup>2</sup> | 2mA | 0.080 | 30min | 10 | Monotherapy | 30sec |
| Sampaio-Junior et al., 2017 | F3<sup>3</sup> | F4<sup>3</sup> | 25cm<sup>2</sup> | 2mA | 0.080 | 30min | 12 | Augmentation | 30sec |

*Note.* Electrode locations are reported according to the EEG 10/20 system. Current densities are reported in mA/cm<sup>2</sup>. Sham stimulation indicates the duration of time that current was applied for giving an initial sensation of tDCS on the scalp. tDCS = transcranial direct current stimulation. <sup>1</sup>Two sham treatment groups were combined.<sup>2</sup>Patients in sham group also

**Table 4.**
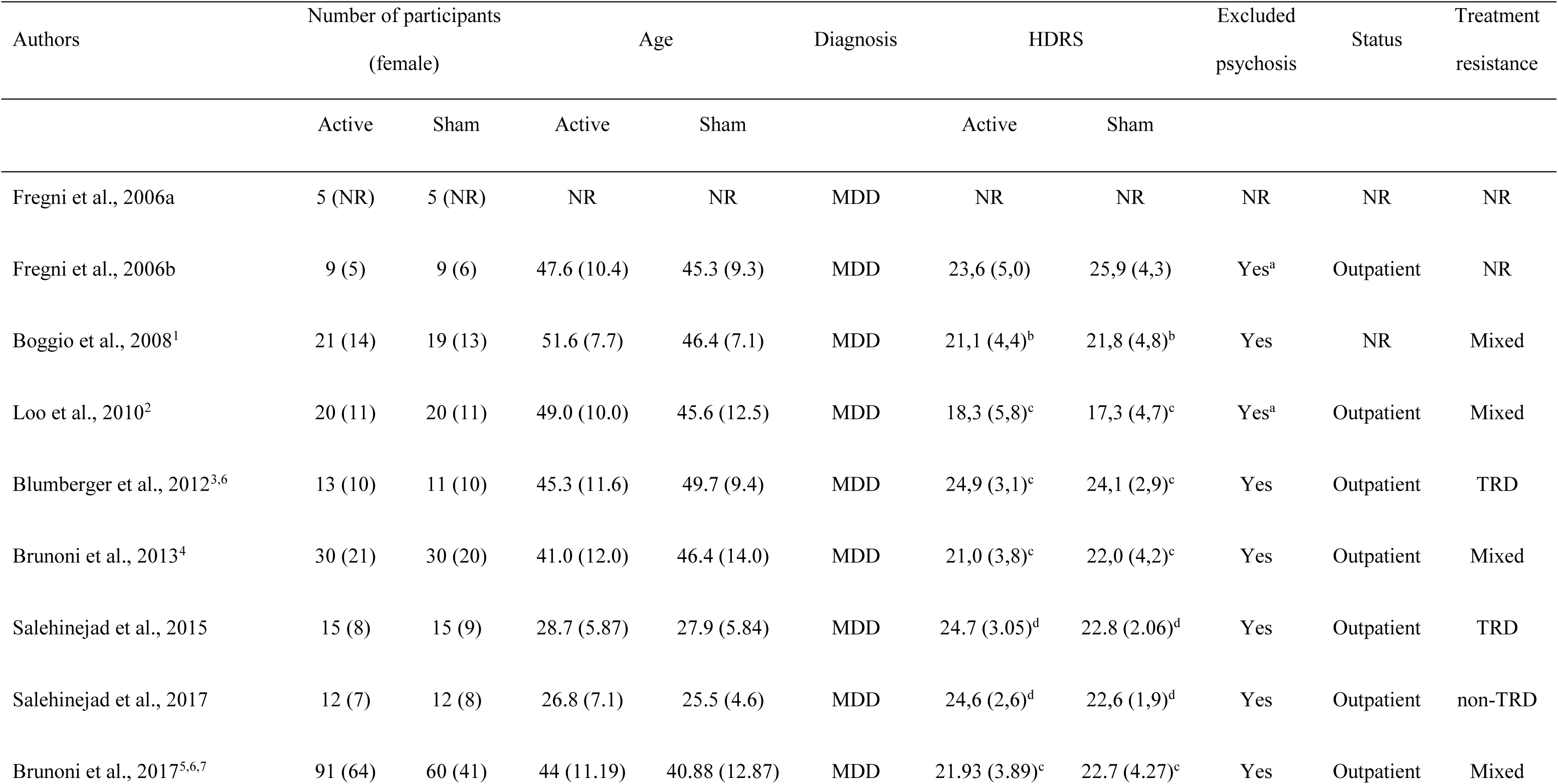
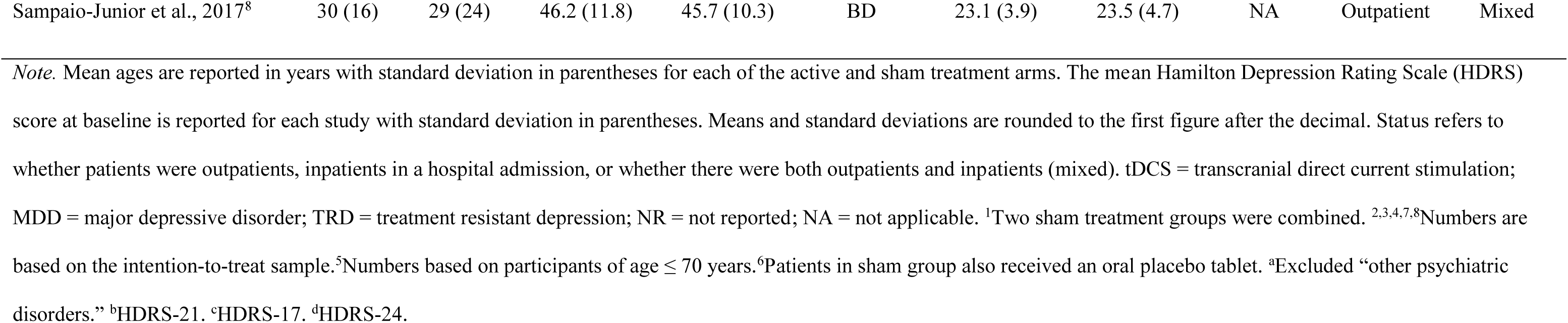
Sample characteristics: tDCS studies

| Authors | Number of participants<br>(female) |  | Age |  | Diagnosis | HDRS |  | Excluded<br>psychosis | Status | Treatment<br>resistance |
| --- | --- | --- | --- | --- | --- | --- | --- | --- | --- | --- |
|  | Active | Sham | Active | Sham |  | Active | Sham |  |  |  |
| Fregni et al., 2006a | 5 (NR) | 5 (NR) | NR | NR | MDD | NR | NR | NR | NR | NR |
| Fregni et al., 2006b | 9 (5) | 9 (6) | 47.6 (10.4) | 45.3 (9.3) | MDD | 23,6 (5,0) | 25,9 (4,3) | Yes<sup>a</sup> | Outpatient | NR |
| Boggio et al., 2008<sup>1</sup> | 21 (14) | 19 (13) | 51.6 (7.7) | 46.4 (7.1) | MDD | 21,1 (4,4)<sup>b</sup> | 21,8 (4,8)<sup>b</sup> | Yes | NR | Mixed |
| Loo et al., 2010<sup>2</sup> | 20 (11) | 20 (11) | 49.0 (10.0) | 45.6 (12.5) | MDD | 18,3 (5,8)<sup>c</sup> | 17,3 (4,7)<sup>c</sup> | Yes<sup>a</sup> | Outpatient | Mixed |
| Blumberger et al., 2012<sup>3,6</sup> | 13 (10) | 11 (10) | 45.3 (11.6) | 49.7 (9.4) | MDD | 24,9 (3,1)<sup>c</sup> | 24,1 (2,9)<sup>c</sup> | Yes | Outpatient | TRD |
| Brunoni et al., 2013<sup>4</sup> | 30 (21) | 30 (20) | 41.0 (12.0) | 46.4 (14.0) | MDD | 21,0 (3,8)<sup>c</sup> | 22,0 (4,2)<sup>c</sup> | Yes | Outpatient | Mixed |
| Salehinejad et al., 2015 | 15 (8) | 15 (9) | 28.7 (5.87) | 27.9 (5.84) | MDD | 24.7 (3.05)<sup>d</sup> | 22.8 (2.06)<sup>d</sup> | Yes | Outpatient | TRD |
| Salehinejad et al., 2017 | 12 (7) | 12 (8) | 26.8 (7.1) | 25.5 (4.6) | MDD | 24,6 (2,6)<sup>d</sup> | 22,6 (1,9)<sup>d</sup> | Yes | Outpatient | non-TRD |
| Brunoni et al., 2017<sup>5,6,7</sup> | 91 (64) | 60 (41) | 44 (11.19) | 40.88 (12.87) | MDD | 21.93 (3.89)<sup>c</sup> | 22.7 (4.27)<sup>c</sup> | Yes | Outpatient | Mixed |
| Sampaio-Junior et al., 2017<sup>8</sup> | 30 (16) | 29 (24) | 46.2 (11.8) | 45.7 (10.3) | BD | 23.1 (3.9) | 23.5 (4.7) | NA | Outpatient | Mixed |

*Note.* Mean ages are reported in years with standard deviation in parentheses for each of the active and sham treatment arms. The mean Hamilton Depression Rating Scale (HDRS) score at baseline is reported for each study with standard deviation in parentheses. Means and standard deviations are rounded to the first figure after the decimal. Status refers to whether patients were outpatients, inpatients in a hospital admission, or whether there were both outpatients and inpatients (mixed). tDCS = transcranial direct current stimulation; MDD = major depressive disorder; TRD = treatment resistant depression; NR = not reported; NA = not applicable. <sup>1</sup>Two sham treatment groups were combined. <sup>2,3,4,7,8</sup>Numbers are based on the intention-to-treat sample. <sup>5</sup>Numbers based on participants of age $\leq 70$ years. <sup>6</sup>Patients in sham group also received an oral placebo tablet. <sup>8</sup>Excluded “other psychiatric disorders.” <sup>b</sup>HDRS-21. <sup>c</sup>HDRS-17. <sup>d</sup>HDRS-24.

Visual inspection of the contour-enhanced funnel plots did not suggest small study effects (Figure 2; Supplementary Material 3). However, due to the small number of studies for treatment modalities other than left-sided high-frequency rTMS and tDCS, these need to be interpreted with caution. The results of our risk of bias assessment are presented in Supplementary Material 4.

**Figure 2.**
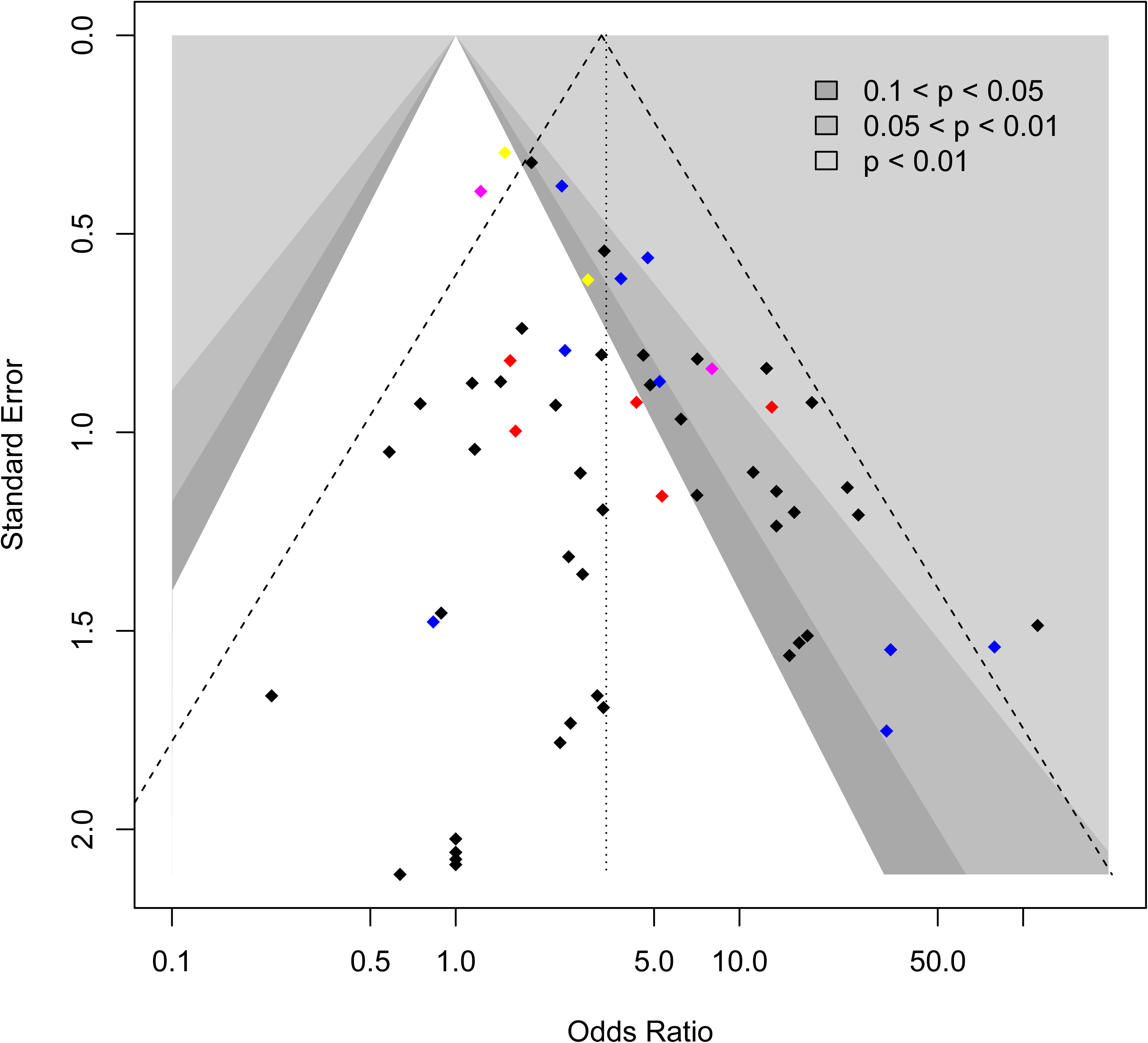
Caption: Contour-enhanced funnel plot of all RCTs included in the meta-analysis of response rates. Legend: rTMS (black); tDCS (navy); TBS (red); dTMS (yellow): sTMS (pink).

### Response and remission rates

Sixty-two comparisons of experimental and sham treatment arms met the inclusion criteria for the meta-analysis of response rates (Table 5; Figure 3), and 50 treatment comparisons for the meta-analysis of remission rates (Table 6; Figure 4).

**Figure 3.**
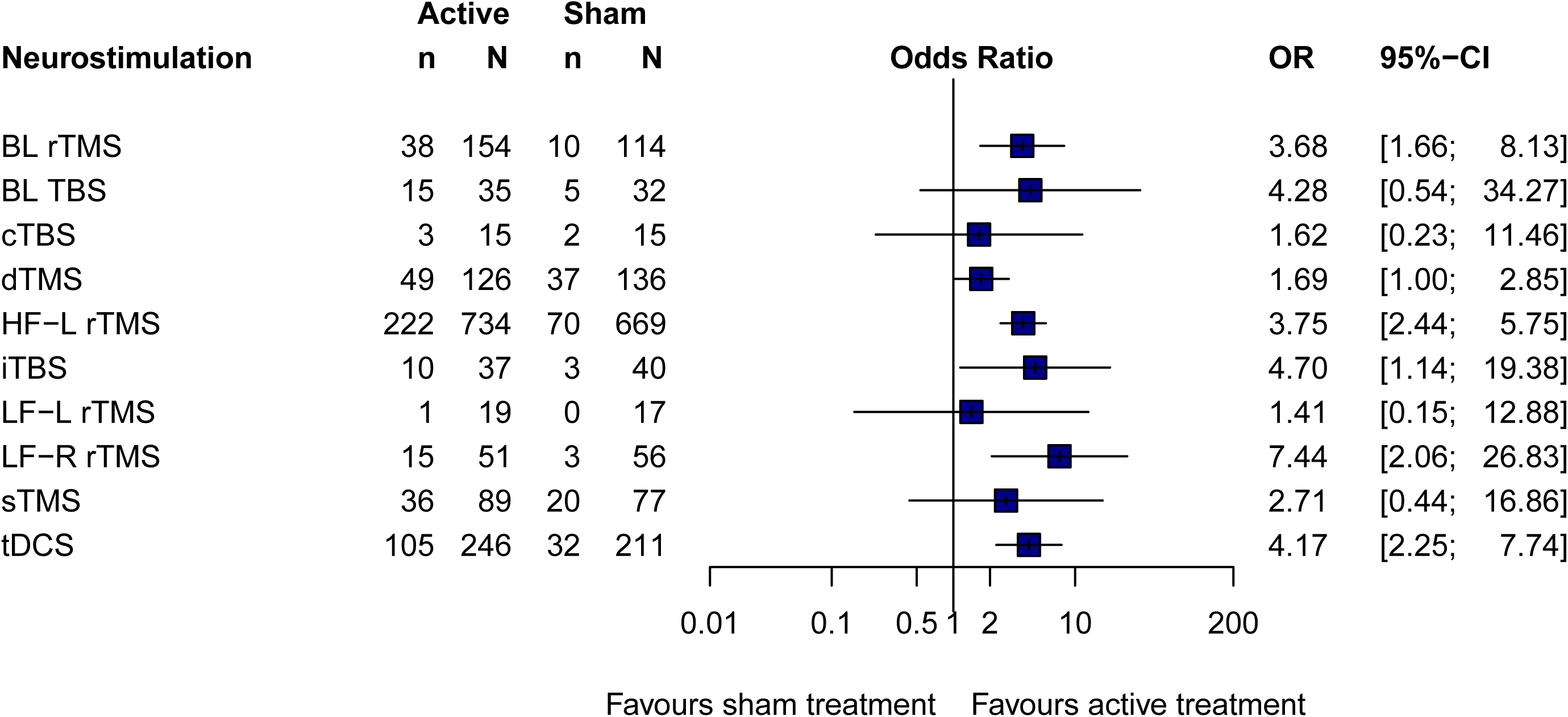
Caption: Forest plot of response rates.

**Figure 4.**
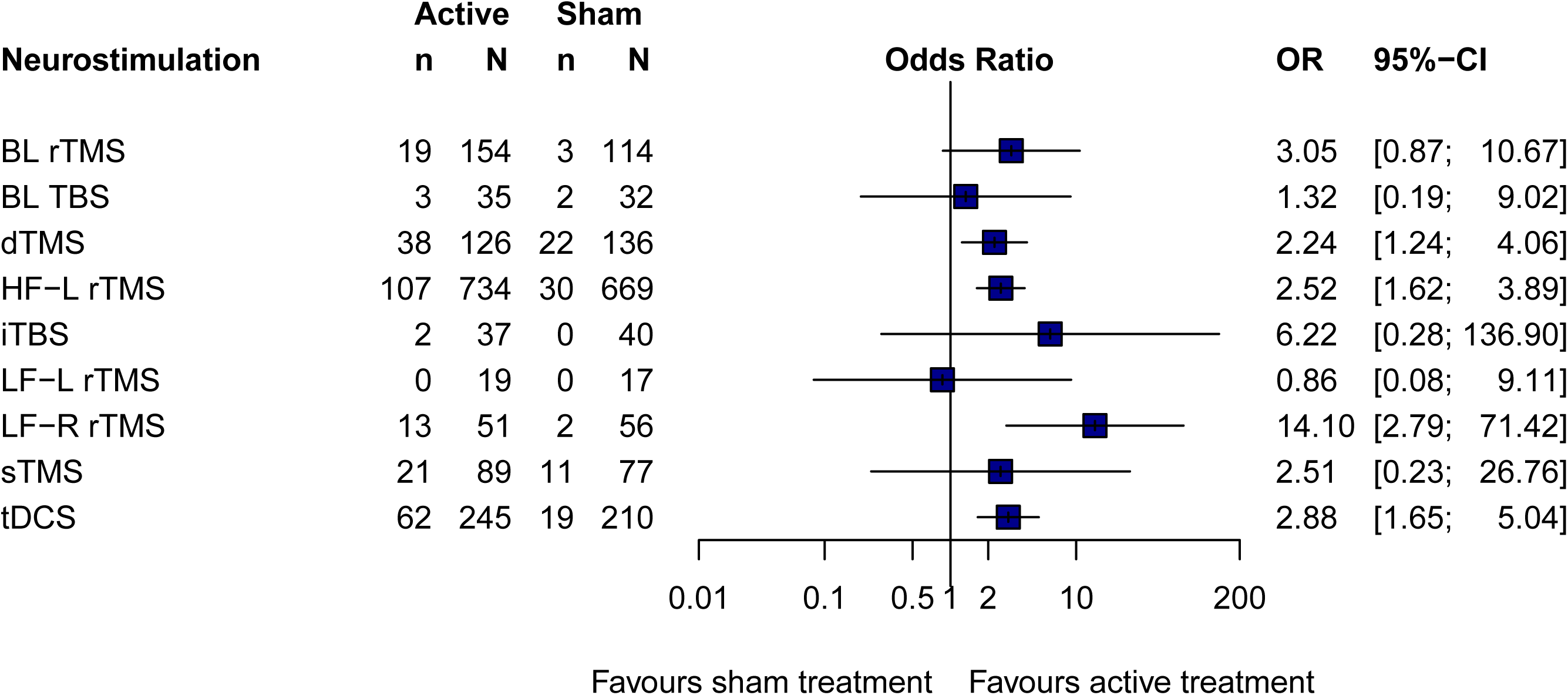
Caption: Forest plot of remission rates.

**Table 5.**
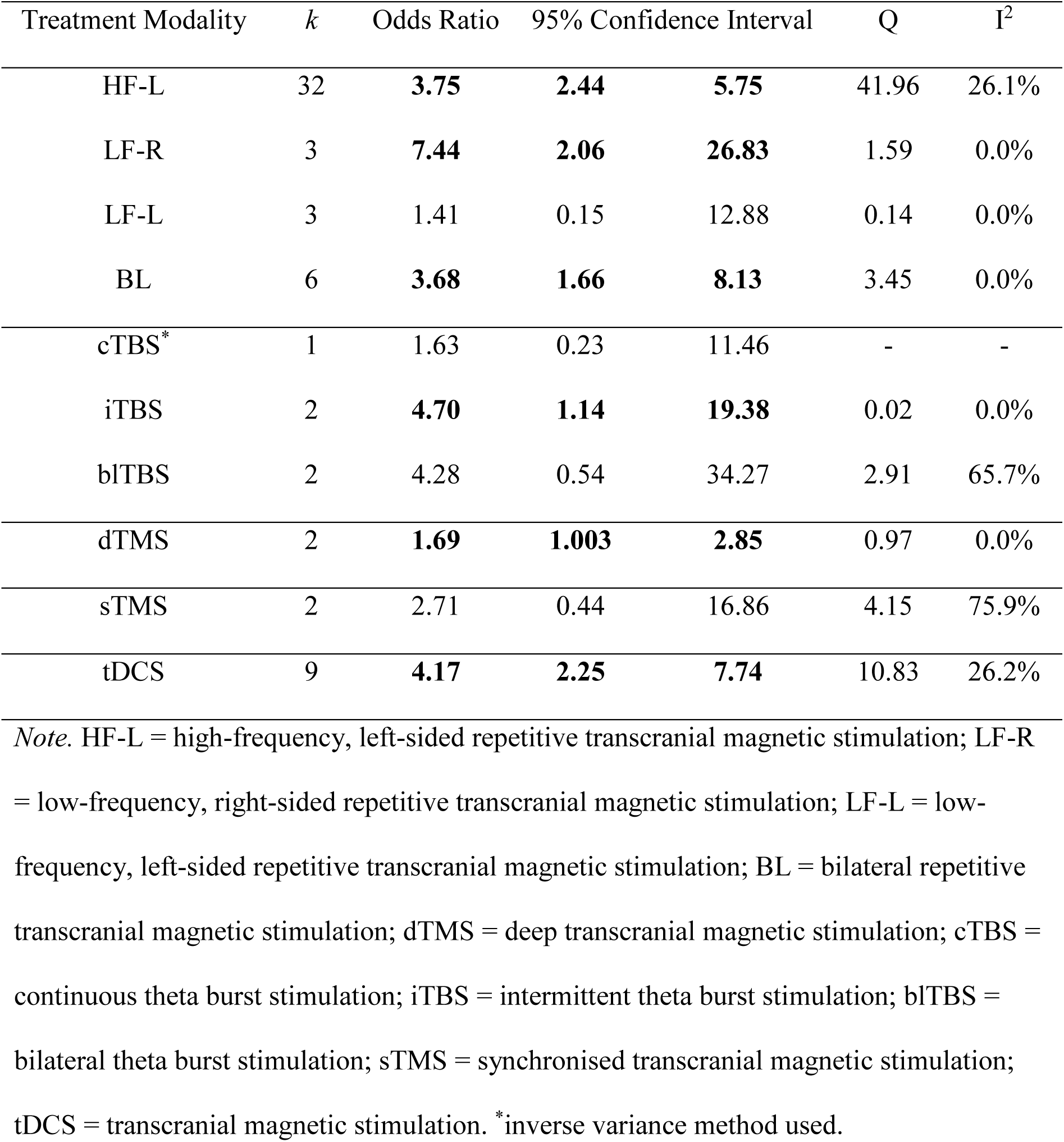
Random-Effects Meta-Analysis of Response Rates

**Table 6.** Random-Effects Meta-Analysis of Remission Rates

*Random-Effects Meta-Analysis of Remission Rates*
| Treatment Modality | *k* | Odds Ratio | 95% Confidence Interval |  |  | Q | I<sup>2</sup> |
| --- | --- | --- | --- | --- | --- | --- | --- |
| HF-L | 26 | **2.52** | **1.62** | **3.89** |  | 25.35 | 1.4% |
| LF-R | 2 | **14.10** | **2.79** | **71.42** |  | 0.50 | 0.0% |
| LF-L | 3 | 0.86 | 0.08 | 9.11 |  | 0.03 | 0.0% |
| BL | 5 | **3.05** | **0.87** | **10.67** |  | 4.48 | 10.7% |
| cTBS | - | - | - | - |  | - | - |
| iTBS* | 1 | 6.22 | 0.28 | 136.90 |  | - | - |
| bITBS* | 1 | 1.32 | 0.19 | 9.02 |  | - | - |
| dTMS | 2 | **2.24** | **1.24** | **4.06** |  | 0.02 | 0.0% |
| sTMS | 2 | 2.51 | 0.23 | 26.76 |  | 4.12 | 75.7% |
| tDCS | 8 | **2.88** | **1.65** | **5.04** |  | 6.32 | 0.0% |

*Note.* HF-L = high-frequency, left-sided repetitive transcranial magnetic stimulation; LF-R = low-frequency, right-sided repetitive transcranial magnetic stimulation; LF-L = low-frequency, left-sided repetitive transcranial magnetic stimulation; BL = bilateral repetitive transcranial magnetic stimulation; dTMS = deep transcranial magnetic stimulation; cTBS = continuous theta burst stimulation; iTBS = intermittent theta burst stimulation; bITBS = bilateral theta burst stimulation; sTMS = synchronised transcranial magnetic stimulation; tDCS = transcranial magnetic stimulation. \*inverse variance method used.

High-frequency rTMS over the left DLPFC (lDLPFC) was associated with improved rates of response as well as remission in comparison with sham treatment. The odds ratio of response was OR = 3.75 compared to sham (*k* = 32, 95% CI [2.44; 5.75]). There was little evidence that the heterogeneity between trials exceeded that expected by chance (I^2^ = 26.1%; Q31= 41.96, *p* = .09). Sensitivity analyses suggested similar effect sizes in trials that had recruited patients with unipolar depression only and those that had recruited both patients with unipolar and bipolar depression (Supplementary Figure 3a).Only one pilot study^38^ had recruited patients with bipolar depression only, but provided no support for antidepressant efficacy (OR = 1.14, 95% CI [0.21; 6.37]). Response rates were greater in trials that (i) excluded patients with psychotic features, (ii) recruited outpatients only, and (iii) recruited either treatment resistant patients only or both treatment resistant patients and those that were not treatment resistant (Supplementary Figures 3b-3d).

The odds of achieving remission were over twice that of sham (*k* = 26, OR = 2.51, 95% CI [1.62; 3.89]). There was no evidence for significant heterogeneity (I^2^ = 1.4%; Q_25_ = 22.35, *p* = .44). Sensitivity analyses for remission rates were in line with those for response rates, although we did not find left-sided high-frequency rTMS to be effective in samples that had recruited both treatment resistant and non-treatment resistant patients (Supplementary Figures 6a-6d).

Low-frequency rTMS over the rDLPFC was also associated with significantly greater response and remission rates than sham stimulation. There was a sevenfold improvement in response rates compared to sham (*k* = 3, OR= 7.44 (95% CI [2.06; 26.83]), with no indication for significant heterogeneity between trials (I^2^ = 0.0%; Q_2_= 1.59, *p* = .45). No sensitivity analyses were conducted due to the small number of treatment comparisons.

The odds of remission were greater than those of sham (*k* = 2, OR = 14.10 (95% CI [2.79; 71.42]). Heterogeneity between trials was not greater than expected due to sampling error (I^2^ = 0.0%; Q_1_ = 0.50, *p* = .48). No sensitivity analyses were conducted due to the small number of treatment comparisons.

Low-frequency rTMS over the lDLPFC was not associated with any significant improvements in rates of response or remission. There were no significant differences in response rates compared to sham (*k* = 3, OR = 1.41, 95% CI [0.15; 12.88]). The heterogeneity between trials did not exceed that expected by chance (I^2^ = 0.0%; Q_2_ = 0.14, *p* = .93), and no sensitivity analyses were conducted due to the small number of treatment comparisons. There were no significant differences in remission rates compared to sham (*k* = 3, OR = 0.86, 95% CI [0.08; 9.11]). The variance in effect sizes between trials was no greater than expected due to sampling error (I^2^ = 0.0%; Q_2_ = 0.03, *p* = .98). No sensitivity analyses were conducted due to the small number of treatment comparisons.

Bilateral rTMS was associated with significant improvement in response but not remission rates compared to sham. There was a significant improvement in response rates compared to sham (*k* = 6, OR = 3.68 (95% CI [1.66; 8.13]), and the variance in effect sizes between trials did not exceed that expected due to sampling error (I^2^ = 0.0%; Q_5_ = 3.45, *p* = .63). Sensitivity analyses suggested subgroup differences according to whether trials had excluded psychotic patients or had recruited patients with diagnosis of MDD only, bipolar depression only, or both MDD and bipolar depression (Supplementary Figures 4a,4b). We found no evidence for a significant improvement in rates of remission associated with bilateral TMS compared to sham (*k* = 5, OR = 3.05, 95% CI [0.87; 10.67]). There was no evidence for significant heterogeneity between trials (I^2^ = 10.7%; Q_4_ = 4.48, *p* = .34), and sensitivity analyses suggested no differences according to any patient characteristics tested (Supplementary Figures 7a,7b).

There were significant improvements in both response and remission rates for dTMS compared to sham. The response rates were marginally higher while statistically significant for dTMS relative to sham (*k* =2, OR = 1.69, 95% CI [1.003; 2.85]). The variance in effect sizes between trials did not exceed that expected due to sampling error (I^2^ = 0.0%; Q_1_ = 0.97, *p* = .33). No sensitivity analyses were conducted due to the small number of treatment comparisons. The remission rates were greater for dTMS compared to sham (*k* = 2, OR = 2.24, 95% CI [1.24; 4.06]). There was no evidence for significant heterogeneity between trials (I^2^ = 0.0%; Q_1_ = 0.02, *p* = 0.88), and no sensitivity analyses were conducted due to the small number of treatment comparisons.

Neither response nor remission rates for sTMS were significantly higher than for sham. There was no evidence for increased response rates compared to sham (*k* = 2, OR = 2.71, 95% CI [0.44; 16.86]). There was significant heterogeneity between these two studies (I^2^ = 75.9%; Q_1_ = 4.15, *p* = .04). No sensitivity analyses were conducted due to the small number of treatment comparisons. There were also no significant improvements in remission rates for sTMS compared to sham (*k*= 2, OR = 2.51 (95% CI [0.23; 26.76]). There was evidence for significant heterogeneity between the two studies though (I^2^ = 75.7%; Q_1_= 4.12, *p* = .04). No sensitivity analyses were conducted due to the small number of treatment comparisons.

iTBS over the lDLPFC was associated with a fivefold improvement in response rates compared to sham (*k* = 2, OR = 4.70 (95% CI [1.14; 19.38]). The heterogeneity between trials did not exceed that expected by chance (I^2^ = 0.0%; Q_1_ = 0.02, *p* = .89). No sensitivity analyses were conducted due to the small number of treatment comparisons. For only one trial^39^ was data on remission rates for iTBS available, with no evidence for antidepressant efficacy compared to sham.

Neither cTBS over the rDLPFC nor bilateral TBS were statistically different from sham in terms of response rates (*k* = 1, OR = 1.63, 95% CI [0.23; 11.46] and *k* = 2, OR = 4.28, 95% CI [0.54; 34.27]). For bilateral TBS there was evidence that the variance in effect sizes between studies was greater than what would be expected due to sampling error (I^2^ = 65.7%; Q_1_ = 2.91, *p* = .09). No sensitivity analyses were conducted due to the small number of treatment comparisons. The only trial of bilateral TBS for which remission rates were available^40^ found no evidence for its antidepressant efficacy compared to sham. No remission rates were available for cTBS.

tDCS was associated with significant improvement in both response and remission rates in comparison to sham stimulation. There was a significant improvement in response rates relative to sham (*k* = 9, OR = 4.17, 95% CI [2.25; 7.74]). There was little evidence for significant heterogeneity between studies (I^2^ = 26.2%; Q_8_ = 10.83, *p* = .21) and sensitivity analyses suggested tDCS to be effective only in patients with non-treatment resistant depression and in trials that had recruited patients with both treatment resistant and non-treatment resistant depression (Supplementary Figure 5).

The analysis of remission rates showed a statistically significant advantage of tDCS compared to sham (*k* = 8, OR = 2.88, 95% CI [1.65; 5.04]). There was no indication for significant heterogeneity between trials (I^2^ = 0.0%; Q_7_ = 6.32, *p* = .50), and sensitivity analyses found that only trials that had recruited patients with both treatment resistant and non-treatment resistant depression provided evidence for antidepressant efficacy (Supplementary Figure 8).

### Effects on continuous measures

Forty-six treatment comparisons reported post-intervention continuous depression scores. There was evidence for the antidepressant efficacy of high-frequency rTMS over the lDLPFC compared to sham (*k* = 29, Hedge’s *g* = -0.72, 95% CI [-0.99; -0.46]), dTMS compared to sham (*k* = 2, Hedge’s *g* = -0.29, 95% CI [-0.55; -0.03]), and tDCS compared to sham (*k* = 7, Hedge’s *g* = -0.76, 95% CI [-1.31; -0.21]). There was evidence for significant heterogeneity between trials for several treatment modalities (Table 7; Figure 5).

**Table 7.**
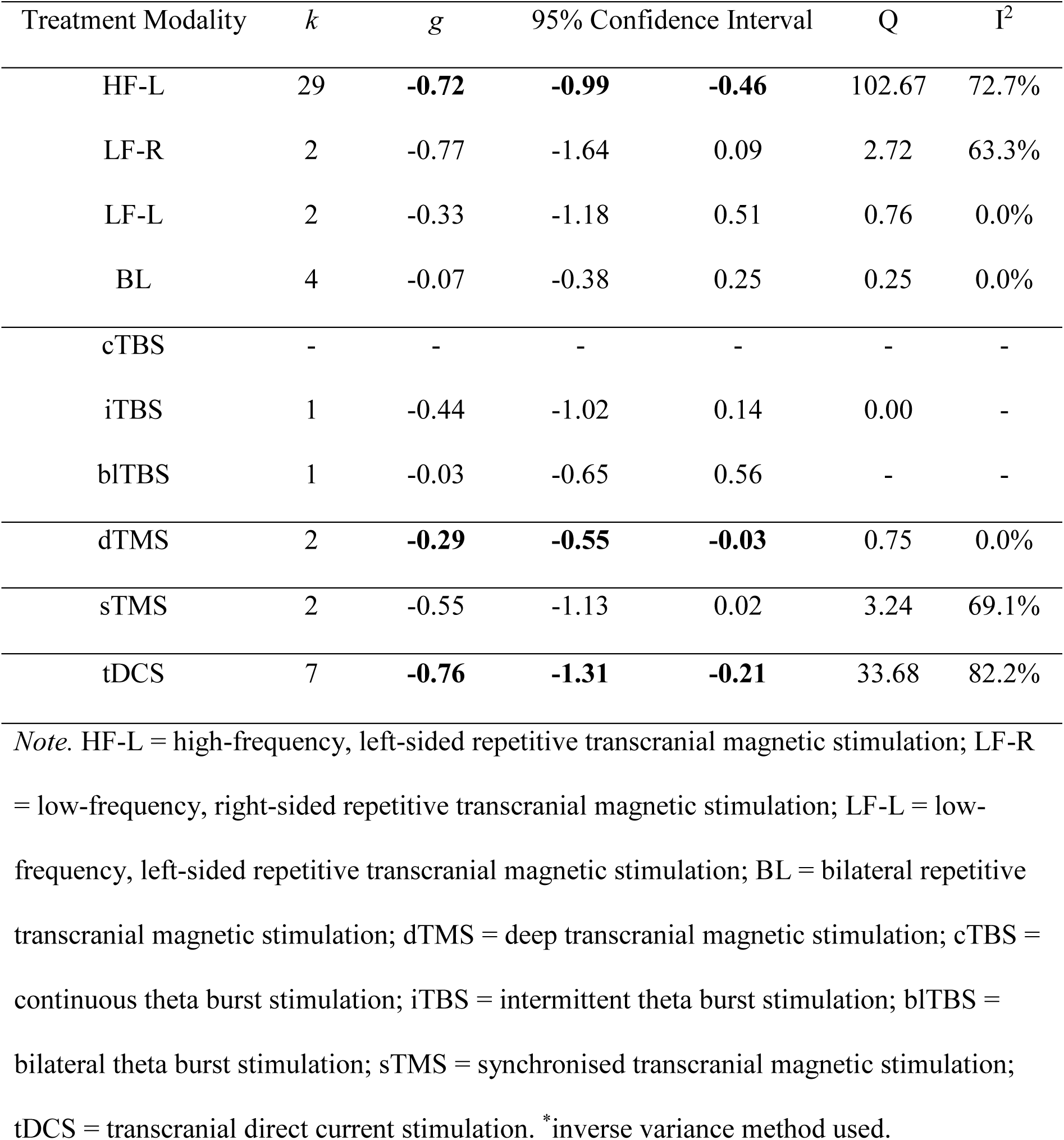
Random-Effects Meta-Analysis of Continuous Treatment Effects

**Figure 5.**
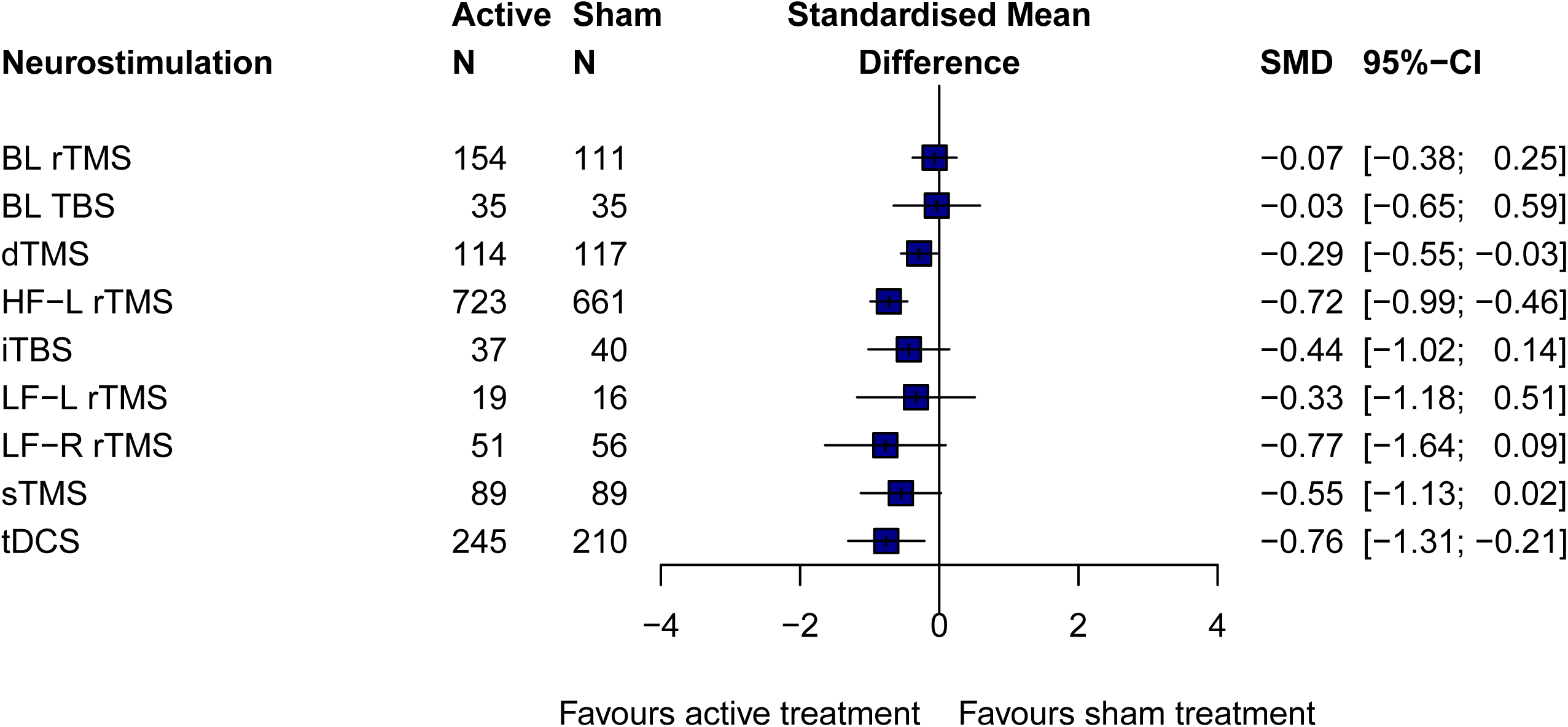
Caption: Forest plot of post-treatment continuous depression scores.

### Acceptability

Sixty-four treatment comparisons were available for all-cause discontinuation rates. There were no significant differences in drop-out rates for any treatment modalities (Table 8; Figure 6).

**Table 8.** Random-Effects Meta-Analysis of All-cause Discontinuation Rates

| Treatment Modality | *k* | Odds Ratio | 95% Confidence Interval |  | Q | I<sup>2</sup> |
| --- | --- | --- | --- | --- | --- | --- |
| HF-L | 35 | 0.86 | 0.60 | 1.23 | 14.58 | 0.0% |
| LF-R | 3 | 0.48 | 0.12 | 1.99 | 0.35 | 0.0% |
| LF-L | 3 | 0.84 | 0.11 | 6.73 | 0.71 | 0.0% |
| BL | 6 | 0.90 | 0.33 | 2.43 | 3.03 | 0.0% |
| cTBS* | 1 | 1.00 | 0.02 | 53.66 | - | - |
| iTBS | 2 | 1.06 | 0.06 | 17.66 | 0.00 | 0.0% |
| BLTBS | 2 | 0.47 | 0.04 | 5.88 | 0.23 | 0.0% |
| dTMS | 2 | 1.03 | 0.32 | 3.36 | 2.10 | 52.3% |
| sTMS | 2 | 0.72 | 0.36 | 1.44 | 0.32 | 0.0% |
| tDCS | 10 | 1.34 | 0.71 | 2.52 | 6.66 | 0.0% |

*Note.* HF-L = high-frequency, left-sided repetitive transcranial magnetic stimulation; LF-R = low-frequency, right-sided repetitive transcranial magnetic stimulation; LF-L = low-frequency, left-sided repetitive transcranial magnetic stimulation; BL = bilateral repetitive transcranial magnetic stimulation; dTMS = deep transcranial magnetic stimulation; cTBS = continuous theta burst stimulation; iTBS = intermittent theta burst stimulation; bITBS = bilateral theta burst stimulation; sTMS = synchronised transcranial magnetic stimulation; tDCS = transcranial direct current stimulation. \*inverse variance method used.

**Figure 6.**
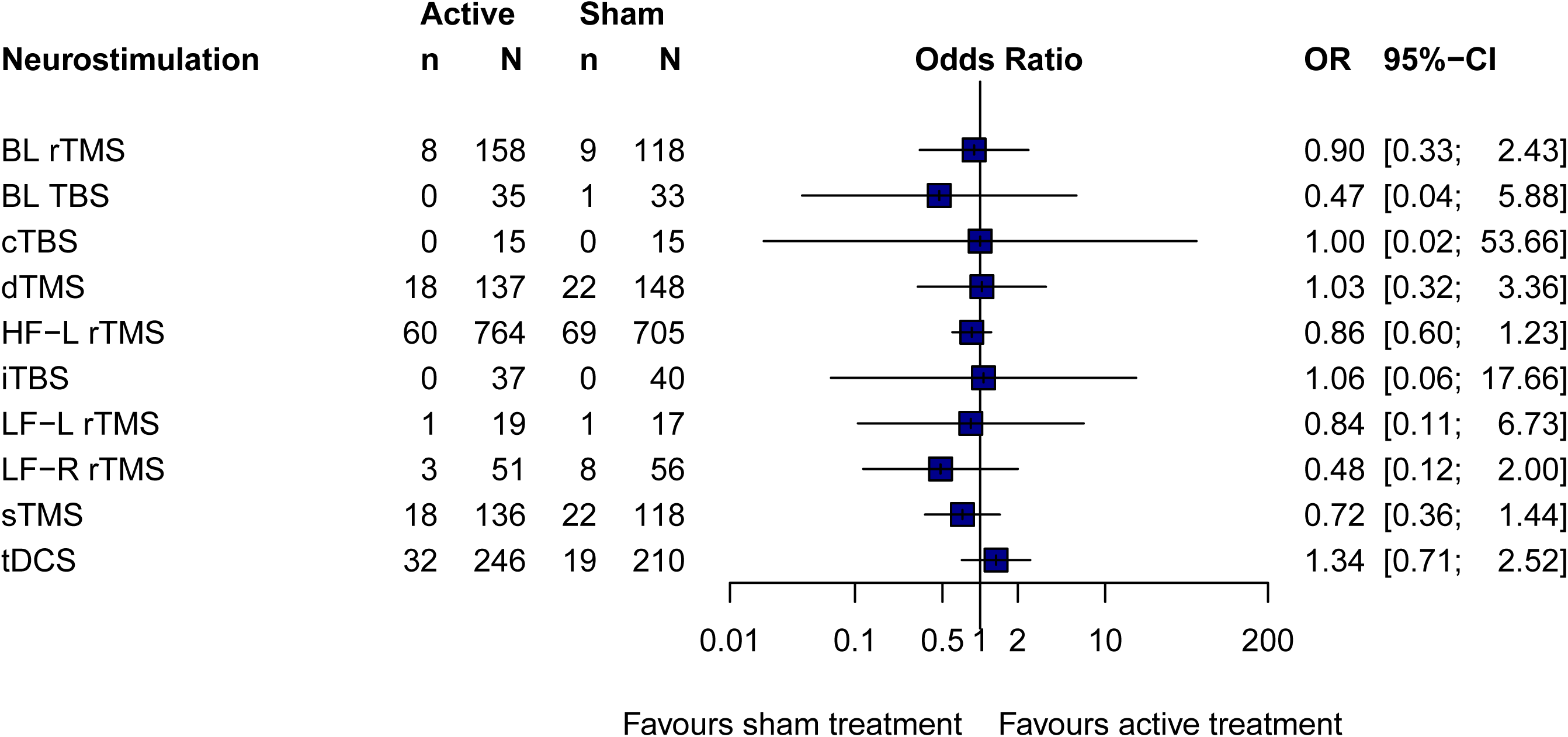
Caption: Forest plot of all-cause discontinuation rates.

## Discussion

The present systematic review and meta-analysis examined the efficacy and acceptability of non-invasive brain stimulation techniques for a current depressive episode in unipolar and bipolar depression. We sought to investigate the efficacy of the brain stimulation techniques without the potential confound of co-initiation of another treatment and in trials which had included randomised allocation to a sham stimulation treatment arm in order to account for potential placebo effects.

The largest evidence base to date is for high-frequency rTMS over the lDLPFC which is associated with 3.75 times greater odds of response than sham stimulation as well as odds of remission that are 2.52 times greater than sham. These findings are consistent with previous systematic reviews and meta-analyses^41^ and have led to the consensus review and treatment guideline by the *Clinical TMS Society* for daily high-frequency rTMS over the lDLPFC for the treatment of medication-resistant or medication-intolerant depressive episodes^42^.

Additional support for treatment efficacy was revealed for low-frequency rTMS over the rDLPFC, which was associated with improved rates of response as well as remission. Bilateral rTMS was associated with higher rates of response but not remission. It is unclear whether any advantages of bilateral rTMS compared to left-sided high-frequency or right-sided low-frequency rTMS would be due to the treatment protocol. As bilateral stimulation delivers a greater number of pulses than unilateral stimulation, unless the number of treatment sessions or the treatment duration are adjusted for accordingly, it is difficult to reliably assess whether the difference in stimulation protocol (bilateral vs. unilateral stimulation) or the difference in the number of stimuli delivered leads to differences in clinical effects^43^.

To date, no studies have directly compared dTMS and standard rTMS protocols. In an exploratory meta-analysis of nine open-label trials, including a total of 150 patients, Kedzior et al.^44^ provided evidence for the antidepressant efficacy of dTMS. The present meta-analysis found that dTMS was associated with 1.69 times greater odds of response and 2.24 greater odds of remission than sham which were statistically significant. While the open-label trials included in Kedzior et al.'s analysis may have overestimated the true efficacy of dTMS, we provide initial support for the clinical efficacy of dTMS that was greater than for sham treatment but less than for high-frequency rTMS over the lDLPFC, low-frequency rTMS over the rDLPFC or bilateral rTMS.

The meta-analytic estimates did not indicate significant treatment effects associated with low-frequency rTMS over the lDLPFC or with sTMS. However, these have been trialled in onlythree^45–47^ and twostudies^21,48^, respectively. Specific treatment effects of TMS that depend on side and frequency of stimulation have been proposed but it may be possible that low-frequency rTMS over the lDLPFC has a marginal effect in at least a small number of patients^47^. Leuchter et al.^48^ found sTMS to only be effective when administered at the individual’s alpha frequency and with a minimum of 80% treatment adherence, suggesting a dose-response relationship.

With theta burst stimulation, the duration of each treatment session is reduced to a few minutes. Our meta-analysis did demonstrate almost five times greater odds of response compared to sham for iTBS over the lDLPFC. However, this estimate is based on two trials only. One trial had examined remission rates as well^39^, reporting remission rates of 0% for sham and 9.1% for active stimulation. The meta-analytic estimates for cTBS and the bilateral modification of TBS did not show any advantage over sham in terms of response rates. The only trial that reported remission rates for bilateral TBS did not provide evidence for its antidepressant efficacy either and no data were available to evaluate remission rates following cTBS.

Transcranial direct current stimulation is a form of neurostimulation that offers greater portability and lower costs relative to TMS. The meta-analysis revealed significant improvements in response and remission rates following tDCS treatment in comparison to sham, which was 4.17 times greater for response rates and 2.88 times greater for remission rates. We have been able to identify the effects of tDCS without potential confounds of co-initiation of another treatment, revealing significantly greater odds of response as well as remission^49^. The clinical efficacy of tDCS is evident also in the non-treatment resistant form of depression, in contrast to most rTMS trials, suggesting that tDCS is a potential initial therapeutic option for depression.

The finding that there were no differences in terms of drop-out rates at study end between the active treatment and sham conditions for any treatment modality suggests that non-invasive brain stimulation is generally well tolerated by patients. We chose all-cause discontinuation rates based on the intention-to-treat sample, representing the most conservative estimate of treatment acceptability.

We chose response and remission rates as our main outcome measures, which are commonly used in the medical sciences and arguably constitute clinically-useful estimates of the antidepressant efficacy of treatment. However, the dichotomisation of outcome data has received criticism because it is known to produce a loss of signal and might inflate Type I error rates, for example an individual who has a 49% reduction in their depressive severity scores would not be included in the clinical response rate while a 51% reduction would be included in the response rate^50^. To address these limitations, we had also analysed continuous depression severity scores. However, outcome data were not reported for each trial, and some missing data could not be obtained. Studies have also suggested that the antidepressant efficacy of active stimulation may separate from sham only after multiple weeks of treatment, for both rTMS^9^ and cTBS^51^. We had examined the acute antidepressant effects at primary study endpoint, and we cannot estimate the long-term effects.

A significant number of TMS studies used active magnetic stimulation with the coil being angulated at 45 or 90 degrees to the scalp surface as sham condition. Because differences in coil orientation may produce considerably different sensations on the scalp and coil angulation might still produce a limited degree of intracortical activity^52^, ensuring a valid control condition constitutes a methodological challenge. One study placed an inactive coil on the patient’s head while discharging an active coil at least one meter away in order to mimic the auditory effects of rTMS^53^.

A more recent approach is to use a specifically designed sham coil that does not generate a magnetic field but is visually and auditorily indistinguishable from an active coil. A meta-analysis by Berlim et al.^54^ found no significant differences between the number of patients who correctly guessed their treatment allocation when comparing active high-frequency left-sided or bilateral rTMS and sham. There were also no significant differences between studies that utilised angulated coils and sham coils. Blinding integrity is less of a methodological hurdle for sTMS trials because neither active stimulation nor sham procedure produce any physical sensation, they look identical, and are comparable in terms of acoustic artefacts. Only few of the more recent modifications of TMS reported on the adequacy of their blinding procedure. Given that cross-over designs are particularly prone to unblinding after cross-over, we included only data corresponding to the initial randomisation in our analyses.

For tDCS, the sham condition typically involves delivering active stimulation for up to 30 seconds, which mimics the initial somatic sensations without inducing a therapeutic effect. However, the adequacy of blinding of tDCS sham has also been called into question^55^.

The clinical trials had enrolled patients based on a diagnostic assessment of clinical symptoms rather than underlying brain pathology. The potential for biological heterogeneity might mask the clinical efficacy of non-invasive brain stimulation in some trials but could not be assessed in the present analysis. We implemented reasonably strict inclusion criteria to limit the influence of a range of potential confounders, for example we excluded RCTs that co-initiated treatment with medication. However, potential effects of specific medications on the clinical efficacy of brain stimulation could not be adequately controlled for as patients often had a large number of heterogeneous treatments prior to enrolling, which might have distorted the clinical effects of brain stimulation.

Finally, compared to the network meta-analysis (NMA) on TMS^29^, we were not able to compare the active treatments. In the NMA priming rTMS seemed most effective. However, the two RCTs that used this treatment modality compared it with another active stimulation and could not be included in the present meta-analysis.

### Conclusion

The present systematic review and meta-analysis supports the efficacy and acceptability of non-invasive brain stimulation techniques in adult unipolar and bipolar depression. The strongest evidence was for high-frequency rTMS over the lDLPFC, followed by low-frequency rTMS over the rDLPFC and bilateral rTMS. Intermittent TBS provides a potential advance in terms of reduced treatment duration and the meta-analysis did find support for improved rates of response. tDCS is a potential treatment for non-resistant depression which has demonstrated efficacy in terms of response as well as remission. All the trials included in the present meta-analysis had included randomised allocation to a sham treatment arm and we had excluded trials in which there was co-initiation of another treatment. Some of the more recent treatment modalities though require additional trials and more direct comparisons between different treatment modalities are warranted.

### Authorship contributions

C.H.Y.F. and J.M. conceived the project; J.M. performed the systematic literature search with supervision by C.H.Y.F; J.M. extracted and analysed the data; D.R.E. reviewed the quality of the extracted data; J.M. wrote the initial draft; C.H.Y.F. critically revised each draft, including interpretation of the data; A.R.B. critically revised the paper. All authors read and approved the final version of this paper. J.M is the guarantor.

### Funding and disclosure

The authors declare no conflict of interest.

## Acknowledgments

We thank Dr Angelo Alonzo & Professor Colleen Loo, Professor Ian Anderson, Dr Lysianne Beynel, Dr Noomane Bouaziz & Dr Dominique Januel, Dr Poul E Buchholtz, Dr Romain Duprat & Professor Chris Baeken, Professor Mark S George, Professor Ehud Klein, Professor Berthold Langguth & Dr Martin Schecklmann, Dr Giuseppe Lanza, Professor Andy Leuchter, Dr Alessandra Minelli, Professor William M McDonald, Professor Declan McLoughlin, Dr Marie-Laure Paillère Martinot, Dr Bill Phillips, Professor Robert M Post, Dr Mohammad Ali Salehinejad, Dr Christos Theleritis, and Professor Abraham Zangen for providing additional data. We are also grateful to those authors who could not provide additional data but responded to our inquiry. The views expressed in this article represent those of the authors and not necessarily those of the individuals who have provided data for the analyses.

## Supplementary material

Supplementary information is available online.

