## Supplementary Materials for "Efficacy and acceptability of non-invasive brain stimulation for the treatment of adult unipolar and bipolar depression: A systematic review and meta-analysis of randomised sham-controlled trials" (from another version of this preprint: version 1, 2018-03-26)

### 1. Search strategy

The following search terms were used: (bipolar disorder OR bipolar depression OR major depression OR unipolar depression OR unipolar disorder) AND (transcranial direct current stimulation OR tDCS OR transcranial magnetic stimulation OR TMS OR theta burst stimulation/ OR TBS OR sTMS OR dTMS), limiting searches to studies in humans and English-language publications. Reference lists of included papers and of recent systematic reviews and meta-analyses (see below) were screened for further studies.

Berlim, M. T., Van den Eynde, F., & Daskalakis, Z. J. (2013a). Clinical utility of transcranial direct current stimulation (tDCS) for treating major depression: a systematic review and meta-analysis of randomized, double-blind and sham-controlled trials. *Journal of psychiatric research, 47*(1), 1-7.

Berlim, M. T., Van den Eynde, F., & Daskalakis, Z. J. (2013b). Clinically meaningful efficacy and acceptability of low-frequency repetitive transcranial magnetic stimulation (rTMS) for treating primary major depression: a meta-analysis of randomized, double-blind and sham-controlled trials. *Neuropsychopharmacology, 38*(4), 543-551.

Berlim, M. T., Van den Eynde, F., Tovar-Perdomo, S., & Daskalakis, Z. (2014). Response, remission and drop-out rates following high-frequency repetitive transcranial magnetic stimulation (rTMS) for treating major depression: a systematic review and meta-analysis of randomized, double-blind and sham-controlled trials. *Psychological Medicine, 44*(02), 225-239.

Brunoni, A. R., Chaimani, A., Moffa, A. H., Razza, L. B., Gattaz, W. F., Daskalakis, Z. J., & Carvalho, A. F. (2017). Repetitive transcranial magnetic stimulation for the acute treatment of major depressive episodes: A systematic review with network meta-analysis. *JAMA psychiatry, 74*(2), 143-152.

Brunoni, A. R., Moffa, A. H., Fregni, F., Palm, U., Padberg, F., Blumberger, D. M., . . . Alonzo, A. (2016). Transcranial direct current stimulation for acute major depressive episodes: meta-analysis of individual patient data. *The British Journal of Psychiatry, 208*(6), 522-531.

Chen, J.-j., Liu, Z., Zhu, D., Li, Q., Zhang, H., Huang, H., . . . Xie, P. (2014). Bilateral vs. unilateral repetitive transcranial magnetic stimulation in treating major depression: a meta-analysis of randomized controlled trials. *Psychiatry Research, 219*(1), 51-57.

Kedzior, K. K., Gellersen, H. M., Brachetti, A. K., & Berlim, M. T. (2015). Deep transcranial magnetic stimulation (DTMS) in the treatment of major depression: an exploratory systematic review and meta-analysis. *Journal of affective disorders, 187*, 73-83.

Lepping, P., Schönfeldt-Lecuona, C., Sambhi, R., Lanka, S., Lane, S., Whittington, R., . . . Poole, R. (2014). A systematic review of the clinical relevance of repetitive transcranial magnetic stimulation. *Acta Psychiatrica Scandinavica, 130*(5), 326-341.

Meron, D., Hedger, N., Garner, M., & Baldwin, D. S. (2015). Transcranial direct current stimulation (tDCS) in the treatment of depression: systematic review and meta-analysis of efficacy and tolerability. *Neuroscience & Biobehavioral Reviews, 57*, 46-62.

Schutter, D. (2009). Antidepressant efficacy of high-frequency transcranial magnetic stimulation over the left dorsolateral prefrontal cortex in double-blind sham-controlled designs: a meta-analysis. *Psychological Medicine, 39*(01), 65-75.

Zhang, Y., Zhu, D., Zhou, X., Liu, Y., Qin, B., Ren, G., & Xie, P. (2015). Bilateral repetitive transcranial magnetic stimulation for treatment-resistant depression: a systematic review and meta-analysis of randomized controlled trials. *Brazilian Journal of Medical and Biological Research, 48*(3), 198-206.

#### 2. Full list of references


Anderson, I. M., Delvai, N. A., Ashim, B., Ashim, S., Lewin, C., Singh, V., . . . Strickland, P. L. (2007). Adjunctive fast repetitive transcranial magnetic stimulation in depression. *The British Journal of Psychiatry, 190*(6), 533-534.

Avery, D. H., Claypoole, K., Robinson, L., Neumaier, J. F., Dunner, D. L., Scheele, L., . . . Roy-Byrne, P. (1999). Repetitive transcranial magnetic stimulation in the treatment of medication-resistant depression: preliminary data. *The Journal of nervous and mental disease, 187*(2), 114-117.

Avery, D. H., Holtzheimer, P. E., Fawaz, W., Russo, J., Neumaier, J., Dunner, D. L., . . . Roy-Byrne, P. (2006). A controlled study of repetitive transcranial magnetic stimulation in medication-resistant major depression. *Biological psychiatry, 59*(2), 187-194.

Baeken, C., Vanderhasselt, M.-A., Remue, J., Herremans, S., Vanderbruggen, N., Zeeuws, D., . . . De Raedt, R. (2013). Intensive HF-rTMS treatment in refractory medication-resistant unipolar depressed patients. *Journal of affective disorders, 151*(2), 625-631.

Bakim, B., Uzun, U. E., Karamustafalioglu, O., Ozcelik, B., Alpak, G., Tankaya, O., . . . Yavuz, B. G. (2012). The combination of antidepressant drug therapy and high-frequency repetitive transcranial magnetic stimulation in medication-resistant depression. *Klinik Psikofarmakoloji Bülteni-Bulletin of Clinical Psychopharmacology, 22*(3), 244-253.

Berman, R. M., Narasimhan, M., Sanacora, G., Miano, A. P., Hoffman, R. E., Hu, X. S., . . . Boutros, N. N. (2000). A randomized clinical trial of repetitive transcranial magnetic stimulation in the treatment of major depression. *Biological psychiatry, 47*(4), 332-337.

Blumberger, D. M., Tran, L. C., Fitzgerald, P. B., Hoy, K. E., & Daskalakis, Z. J. (2012). A randomized double-blind sham-controlled study of transcranial direct current stimulation for treatment-resistant major depression. *Frontiers in psychiatry, 3*.

Boggio, P. S., Rigonatti, S. P., Ribeiro, R. B., Myczkowski, M. L., Nitsche, M. A., Pascual-Leone, A., & Fregni, F. (2008). A randomized, double-blind clinical trial on the efficacy of cortical direct current stimulation for the treatment of major depression. *International Journal of Neuropsychopharmacology, 11*(2), 249-254.

Bortolomasi, M., Minelli, A., Fuggetta, G., Perini, M., Comencini, S., Fiaschi, A., & Manganotti, P. (2007). Long-lasting effects of high frequency repetitive transcranial magnetic stimulation in major depressed patients. *Psychiatry research, 150*(2), 181-186.

Boutros, N. N., Gueorguieva, R., Hoffman, R. E., Oren, D. A., Feingold, A., & Berman, R. M. (2002). Lack of a therapeutic effect of a 2-week sub-threshold transcranial magnetic stimulation course for treatment-resistant depression. *Psychiatry research, 113*(3), 245-254.

Brunoni, A. R., Moffa, A. H., Sampaio-Junior, B., Borrione, L., Moreno, M. L., Fernandes, R. A., . . . Razza, L. B. (2017). trial of Electrical Direct-current Therapy versus Escitalopram for Depression. *New England Journal of Medicine, 376*(26), 2523-2533.

Brunoni, A. R., Valiengo, L., Baccaro, A., Zanão, T. A., de Oliveira, J. F., Goulart, A., . . . Fregni, F. (2013). The sertraline vs electrical current therapy for treating depression clinical study: results from a factorial, randomized, controlled trial. *JAMA psychiatry, 70*(4), 383-391.

Chen, S.-J., Chang, C.-H., Tsai, H.-C., Chen, S.-T., & Lin, C. C. (2013). Superior antidepressant effect occurring 1 month after rTMS: add-on rTMS for subjects with medication-resistant depression. *Neuropsychiatric disease and treatment, 9*, 397.

Concerto, C., Lanza, G., Cantone, M., Ferri, R., Pennisi, G., Bella, R., & Aguglia, E. (2015). Repetitive transcranial magnetic stimulation in patients with drug-resistant major depression: a six-month clinical follow-up study. *International journal of psychiatry in clinical practice, 19*(4), 252-258.

Duprat, R., Desmyter, S., van Heeringen, K., Van den Abbeele, D., Tandt, H., Bakic, J., . . . Van Autreve, S. (2016). Accelerated intermittent theta burst stimulation treatment in medication-resistant major depression: A fast road to remission? *Journal of affective disorders, 200*, 6-14.

Eschweiler, G. W., Wegerer, C., Schlotter, W., Spandl, C., Stevens, A., Bartels, M., & Buchkremer, G. (2000). Left prefrontal activation predicts therapeutic effects of repetitive transcranial magnetic stimulation (rTMS) in major depression. *Psychiatry Research: Neuroimaging, 99*(3), 161-172.

Fitzgerald, P. B., Benitez, J., de Castella, A., Daskalakis, Z. J., Brown, T. L., & Kulkarni, J. (2006). A randomized, controlled trial of sequential bilateral repetitive transcranial magnetic stimulation for treatment-resistant depression. *American Journal of Psychiatry, 163*(1), 88-94.

Fitzgerald, P. B., Brown, T. L., Marston, N. A., Daskalakis, Z. J., de Castella, A., & Kulkarni, J. (2003). Transcranial magnetic stimulation in the treatment of depression: a double-blind, placebo-controlled trial. *Archives of General Psychiatry, 60*(10), 1002-1008.

Fitzgerald, P. B., Hoy, K. E., Elliot, D., McQueen, S., Wambeek, L. E., & Daskalakis, Z. J. (2016). A negative double-blind controlled trial of sequential bilateral rTMS in the treatment of bipolar depression. *Journal of affective disorders, 198*, 158-162.

Fitzgerald, P. B., Hoy, K. E., Herring, S. E., McQueen, S., Peachey, A. V., Segrave, R. A., . . . Daskalakis, Z. J. (2012). A double blind randomized trial of unilateral left and bilateral prefrontal cortex transcranial magnetic stimulation in treatment resistant major depression. *Journal of affective disorders, 139*(2), 193-198.

Fregni, F., Boggio, P. S., Nitsche, M. A., Marcolin, M. A., Rigonatti, S. P., & Pascual‐Leone, A. (2006). Treatment of major depression with transcranial direct current stimulation. *Bipolar disorders, 8*(2), 203-204.

Fregni, F., Boggio, P. S., Nitsche, M. A., Rigonatti, S. P., & Pascual‐Leone, A. (2006). Cognitive effects of repeated sessions of transcranial direct current stimulation in patients with depression. *Depression and anxiety, 23*(8), 482-484.

Garcia-Toro, M., Mayol, A., Arnillas, H., Capllonch, I., Ibarra, O., Crespí, M., . . . Lafuente, L. (2001). Modest adjunctive benefit with transcranial magnetic stimulation in medication-resistant depression. *Journal of affective disorders, 64*(2), 271-275.

George, M. S., Lisanby, S. H., Avery, D., McDonald, W. M., Durkalski, V., Pavlicova, M., . . . Zarkowski, P. (2010). Daily left prefrontal transcranial magnetic stimulation therapy for major depressive disorder: a sham-controlled randomized trial. *Archives of General Psychiatry, 67*(5), 507-516.

George, M. S., Nahas, Z., Molloy, M., Speer, A. M., Oliver, N. C., Li, X.-B., . . . Ballenger, J. C. (2000). A controlled trial of daily left prefrontal cortex TMS for treating depression. *Biological psychiatry, 48*(10), 962-970.

George, M. S., Wassermann, E. M., Kimbrell, T. A., Little, J. T., Williams, W. E., Danielson, A. L., . . . Post, R. M. (1997). Mood improvement following daily left prefrontal repetitive transcranial magnetic stimulation in patients with depression: a placebo-controlled crossover trial. *American Journal of Psychiatry, 154*(12), 1752-1756.

Hansen, P. E. B., Videbech, P., Sturlason, R., Clemmensen, K., Jensen, H. M., & Vestergaard, P. (2004). Repetitive transcranial magnetic stimulation as add-on antidepressant treatment. The applicability of the method in a clinical setting. *Nordic journal of psychiatry, 58*(6), 455-457.

Hernández-Ribas, R., Deus, J., Pujol, J., Segalàs, C., Vallejo, J., Menchón, J. M., . . . Soriano-Mas, C. (2013). Identifying brain imaging correlates of clinical response to repetitive transcranial magnetic stimulation (rTMS) in major depression. *Brain stimulation, 6*(1), 54-61.

Holtzheimer, P. E., Russo, J., Claypoole, K. H., Roy‐Byrne, P., & Avery, D. H. (2004). Shorter duration of depressive episode may predict response to repetitive transcranial magnetic stimulation. *Depression and anxiety, 19*(1), 24-30.

Jakob, F., Brakemeier, E.-L., Schommer, N. C., Quante, A., Merkl, A., Danker-Hopfe, H., . . . Bajbouj, M. (2008). Ultrahigh frequency repetitive transcranial magnetic stimulation in unipolar depression. *Journal of clinical psychopharmacology, 28*(4), 474-476.

Januel, D., Dumortier, G., Verdon, C.-M., Stamatiadis, L., Saba, G., Cabaret, W., . . . Kalalou, K. (2006). A double-blind sham controlled study of right prefrontal repetitive transcranial magnetic stimulation (rTMS): therapeutic and cognitive effect in medication free unipolar depression during 4 weeks. *Progress in Neuro-Psychopharmacology and Biological Psychiatry, 30*(1), 126-130.

Jin, Y., & Phillips, B. (2014). A pilot study of the use of EEG-based synchronized Transcranial Magnetic Stimulation (sTMS) for treatment of Major Depression. *BMC psychiatry, 14*(1), 13.

Kimbrell, T. A., Little, J. T., Dunn, R. T., Frye, M. A., Greenberg, B. D., Wassermann, E. M., . . . Benson, B. E. (1999). Frequency dependence of antidepressant response to left prefrontal repetitive transcranial magnetic stimulation (rTMS) as a function of baseline cerebral glucose metabolism. *Biological psychiatry, 46*(12), 1603-1613.

Kreuzer, P. M., Schecklmann, M., Lehner, A., Wetter, T. C., Poeppl, T. B., Rupprecht, R., . . . Langguth, B. (2015). The ACDC pilot trial: targeting the anterior cingulate by double cone coil rTMS for the treatment of depression. *Brain stimulation, 8*(2), 240-246.

Leuchter, A. F., Cook, I. A., Feifel, D., Goethe, J. W., Husain, M., Carpenter, L. L., . . . Bhati, M. T. (2015). Efficacy and safety of low-field synchronized transcranial magnetic stimulation (sTMS) for treatment of major depression. *Brain stimulation, 8*(4), 787-794.

Levkovitz, Y., Isserles, M., Padberg, F., Lisanby, S. H., Bystritsky, A., Xia, G., . . . Dannon, P. (2015). Efficacy and safety of deep transcranial magnetic stimulation for major depression: a prospective multicenter randomized controlled trial. *World Psychiatry, 14*(1), 64-73.

Li, C.-T., Chen, M.-H., Juan, C.-H., Huang, H.-H., Chen, L.-F., Hsieh, J.-C., . . . Lee, Y.-C. (2014). Efficacy of prefrontal theta-burst stimulation in refractory depression: a randomized sham-controlled study. *Brain, 137*(7), 2088-2098.

Lingeswaran, A. (2011). Repetitive transcranial magnetic stimulation in the treatment of depression: A randomized, double-blind, placebo-controlled trial. *Indian journal of psychological medicine, 33*(1), 35.

Loo, C., Mitchell, P., Sachdev, P., McDarmont, B., Parker, G., & Gandevia, S. (1999). Double-blind controlled investigation of transcranial magnetic stimulation for the treatment of resistant major depression. *American Journal of Psychiatry, 156*(6), 946-948.

Loo, C. K., Sachdev, P., Martin, D., Pigot, M., Alonzo, A., Malhi, G. S., . . . Mitchell, P. (2010). A double-blind, sham-controlled trial of transcranial direct current stimulation for the treatment of depression. *International Journal of Neuropsychopharmacology, 13*(1), 61-69.

McDonald, W. M., Easley, K., Byrd, E. H., Holtzheimer, P., Tuohy, S., Woodard, J. L., . . . Epstein, C. M. (2006). Combination rapid transcranial magnetic stimulation in treatment refractory depression. *Neuropsychiatric disease and treatment, 2*(1), 85.

Nahas, Z., Kozel, F. A., Li, X., Anderson, B., & George, M. S. (2003). Left prefrontal transcranial magnetic stimulation (TMS) treatment of depression in bipolar affective disorder: a pilot study of acute safety and efficacy. *Bipolar disorders, 5*(1), 40-47.

O'Reardon, J. P., Solvason, H. B., Janicak, P. G., Sampson, S., Isenberg, K. E., Nahas, Z., . . . Loo, C. (2007). Efficacy and safety of transcranial magnetic stimulation in the acute treatment of major depression: a multisite randomized controlled trial. *Biological psychiatry, 62*(11), 1208-1216.

Padberg, F., Zwanzger, P., Thoma, H., Kathmann, N., Haag, C., Greenberg, B. D., . . . Möller, H.-J. (1999). Repetitive transcranial magnetic stimulation (rTMS) in pharmacotherapy-refractory major depression: comparative study of fast, slow and sham rTMS. *Psychiatry research, 88*(3), 163-171.

Paillère Martinot, M.-L., Galinowski, A., Ringuenet, D., Gallarda, T., Lefaucheur, J.-P., Bellivier, F., . . . Martinot, J.-L. (2010). Influence of prefrontal target region on the efficacy of repetitive transcranial magnetic stimulation in patients with medication-resistant depression: a [18F]-fluorodeoxyglucose PET and MRI study. *International Journal of Neuropsychopharmacology, 13*(1), 45-59.

Pallanti, S., Bernardi, S., Di Rollo, A., Antonini, S., & Quercioli, L. (2010). Unilateral low frequency versus sequential bilateral repetitive transcranial magnetic stimulation: is simpler better for treatment of resistant depression? *Neuroscience, 167*(2), 323-328.

Prasser, J., Schecklmann, M., Poeppl, T. B., Frank, E., Kreuzer, P. M., Hajak, G., . . . Langguth, B. (2015). Bilateral prefrontal rTMS and theta burst TMS as an add-on treatment for depression: a randomized placebo controlled trial. *The World Journal of Biological Psychiatry, 16*(1), 57-65.

Salehinejad, M. A., Ghanavai, E., Rostami, R., & Nejati, V. (2017). Cognitive control dysfunction in emotion dysregulation and psychopathology of major depression (MD): Evidence from transcranial brain stimulation of the dorsolateral prefrontal cortex (DLPFC). *Journal of affective disorders, 210*, 241-248.

Salehinejad, M. A., Rostami, R., & Ghanavati, E. (2015). Transcranial direct current stimulation of dorsolateral prefrontal cortex of major depression: Improving visual working memory, reducing depressive symptoms. *NeuroRegulation, 2*(1), 37-49.

Speer, A. M., Wassermann, E. M., Benson, B. E., Herscovitch, P., & Post, R. M. (2014). Antidepressant efficacy of high and low frequency rTMS at 110% of motor threshold versus sham stimulation over left prefrontal cortex. *Brain stimulation, 7*(1), 36-41.

Su, T.-P., Huang, C.-C., & Wei, I.-H. (2005). Add-on rTMS for medication-resistant depression: a randomized, double-blind, sham-controlled trial in Chinese patients. *The Journal of clinical psychiatry, 66*(7), 930-937.

Tavares, D. F., Myczkowski, M. L., Alberto, R. L., Valiengo, L., Rios, R. M., Gordon, P., . . . Marcolin, M. A. (2017). Treatment of Bipolar Depression with Deep TMS (dTMS): Results from a Double-Blind, Randomized, Parallel Group, Sham-Controlled Clinical Trial. *Neuropsychopharmacology*.

Theleritis, C., Sakkas, P., Paparrigopoulos, T., Vitoratou, S., Tzavara, C., Bonaccorso, S., . . . Psarros, C. (2017). Two Versus One High-Frequency Repetitive Transcranial Magnetic Stimulation Session per Day for Treatment-Resistant Depression: A Randomized Sham-Controlled Trial. *The journal of ECT, 33*(3), 190-197.

Zheng, H., Zhang, L., Li, L., Liu, P., Gao, J., Liu, X., . . . Zhang, Z. (2010). High-frequency rTMS treatment increases left prefrontal myo-inositol in young patients with treatment-resistant depression. *Progress in Neuro-Psychopharmacology and Biological Psychiatry, 34*(7), 1189-1195.

#### 3. Small study effects

Supplementary Figure 1. Contour-enhanced funnel plot of all RCTs included in the meta-analysis of remission rates.

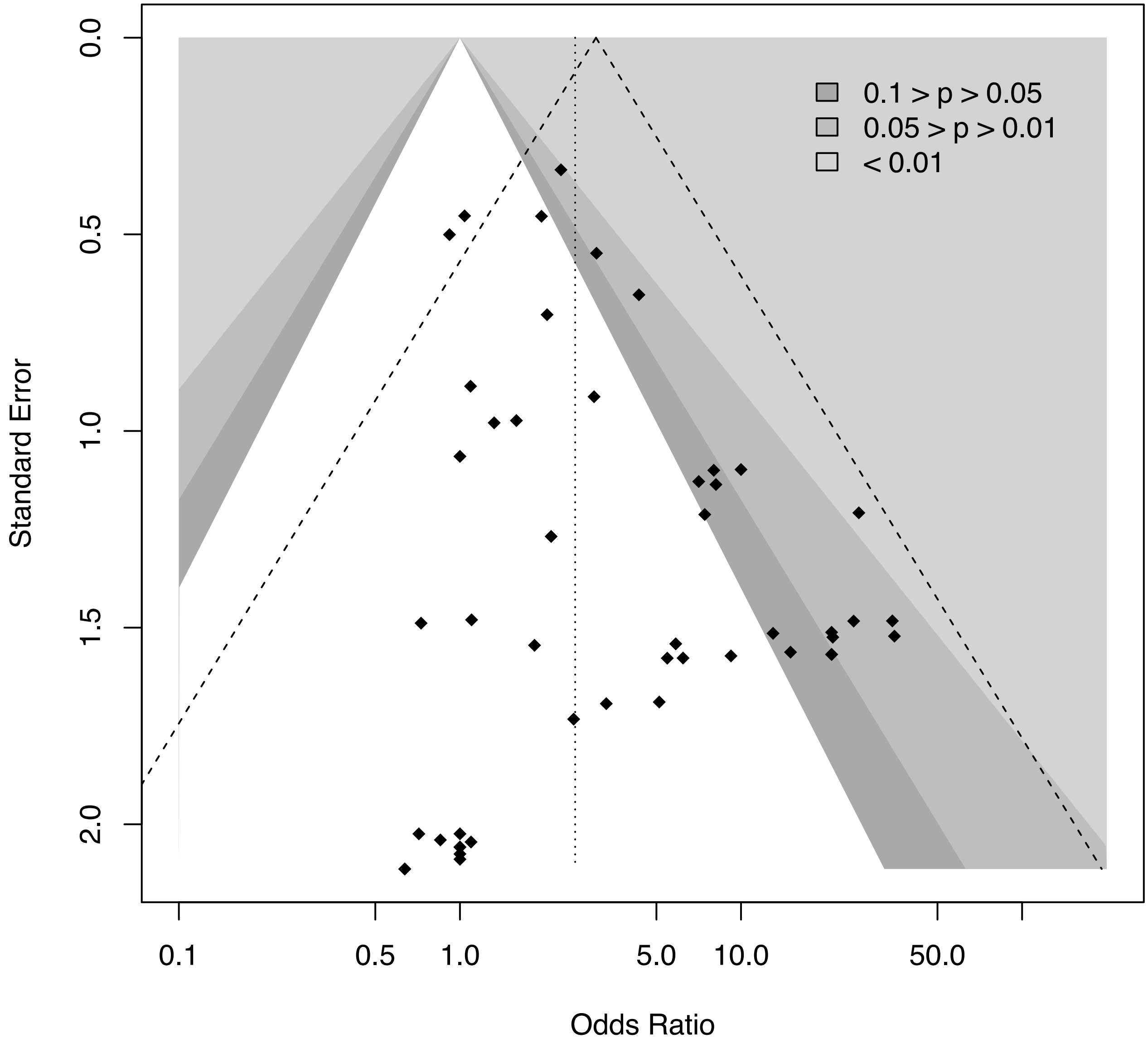

Supplementary Figure 2. Contour-enhanced funnel plot of all RCTs included in the meta-analysis of post-treatment continuous depression scores.

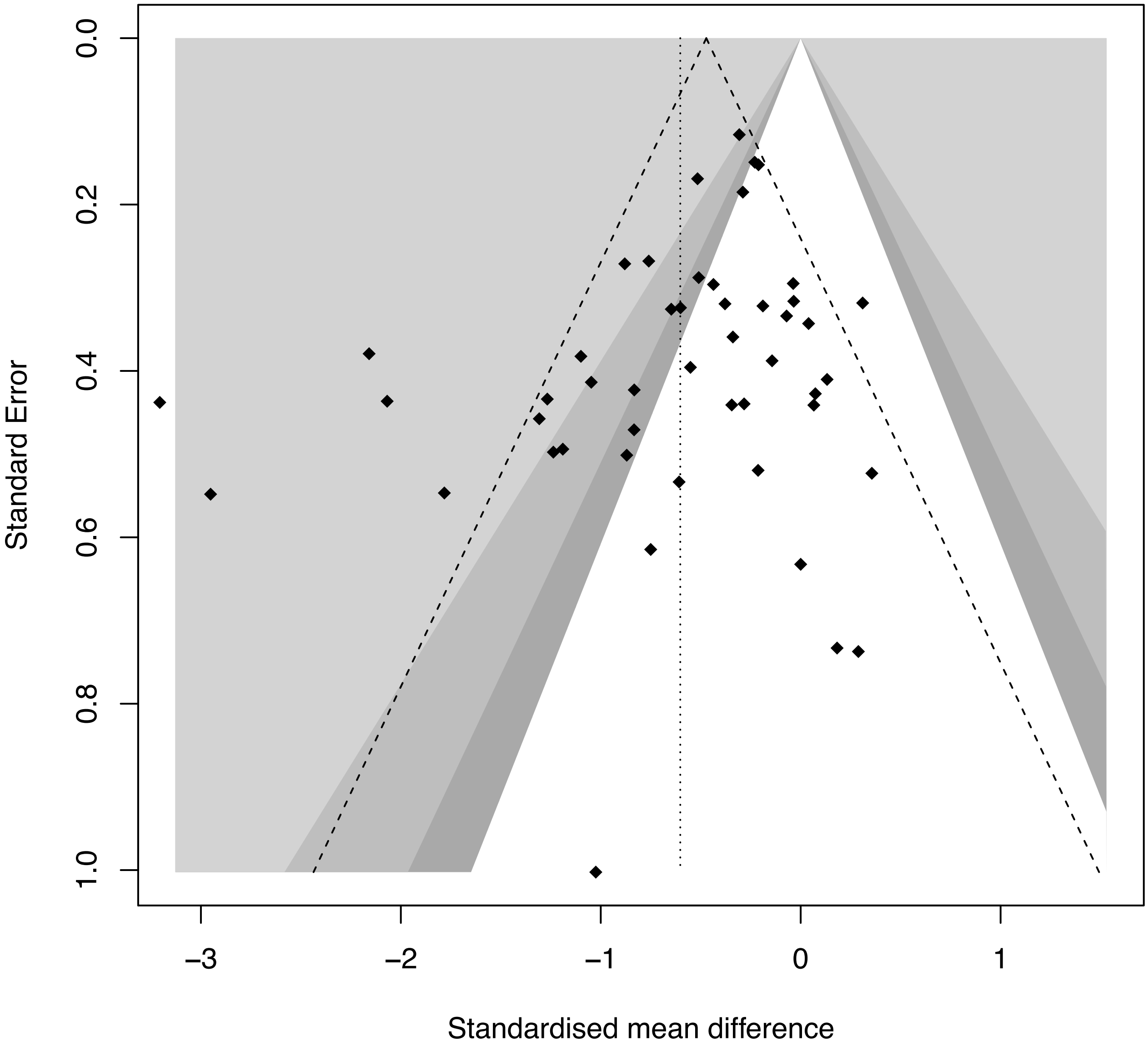

### 4. Sensitivity analyses – response rates.

Supplementary Figure 3a. Forest plot of HF-L (diagnosis).

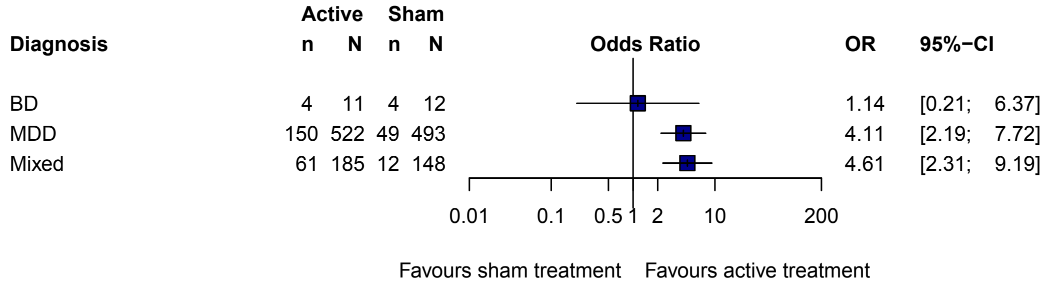


Supplementary Figure 3b. Forest plot of HF-L (exclusion psychosis).

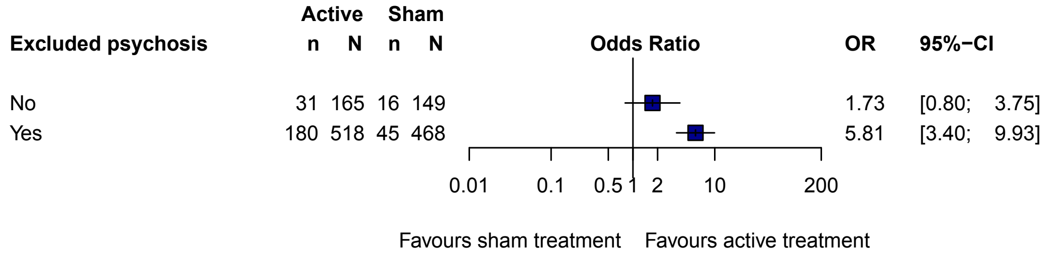


Supplementary Figure 3c. Forest plot of HF-L (hospitalisation status).

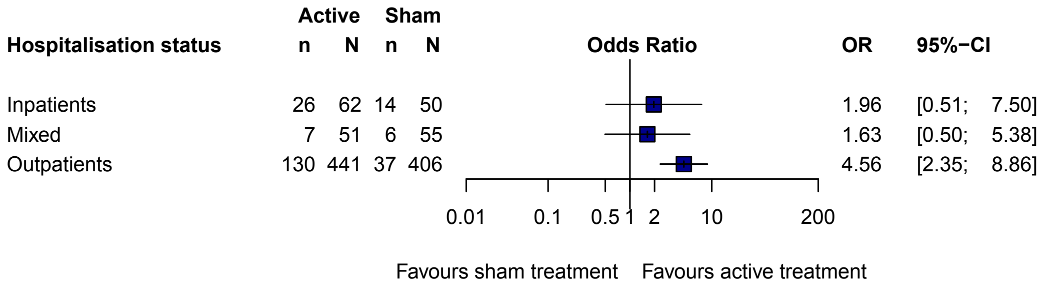


Supplementary Figure 3d. Forest plot of HF-L (treatment resistance).

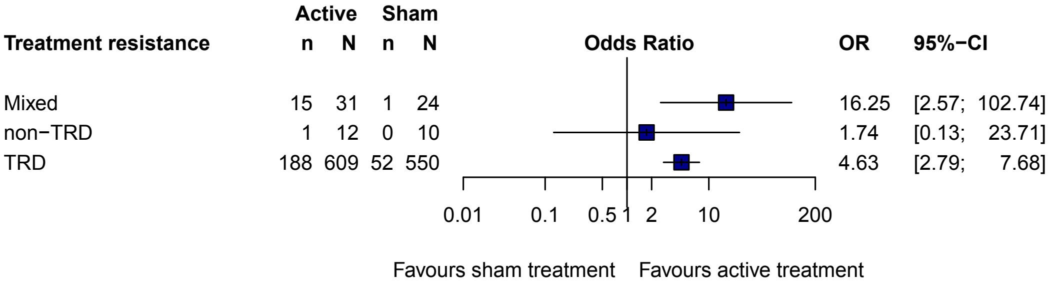

Supplementary Figure 4a. Forest plot of BL (diagnosis).

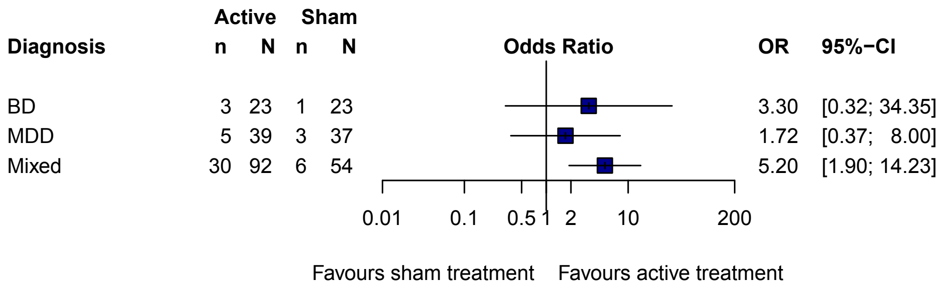


Supplementary Figure 4b. Forest plot of BL (exclusion psychosis).

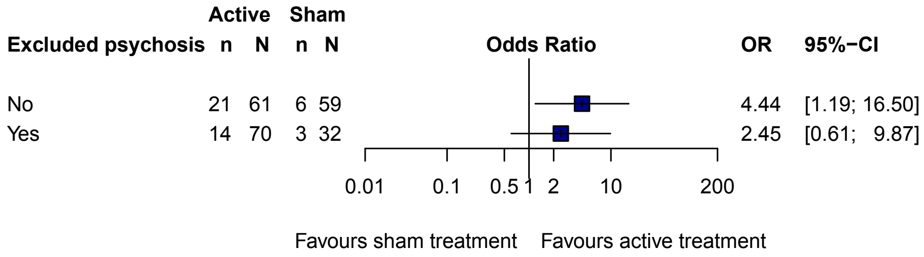


Supplementary Figure 5. Forest plot of tDCS (treatment resistance).

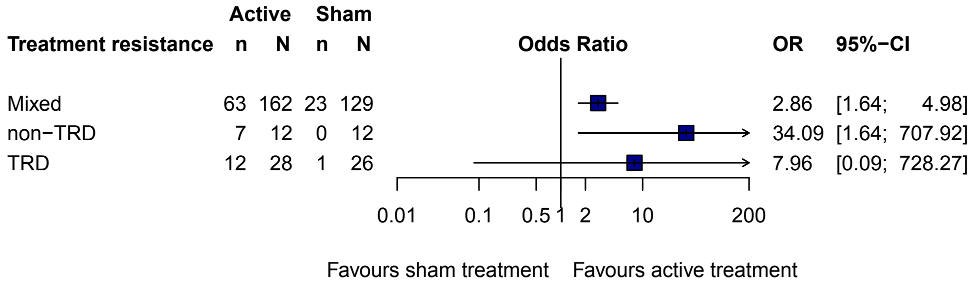

### 5. Sensitivity analyses – remission rates.

Supplementary Figure 6a. Forest plot of HF-L (diagnosis).

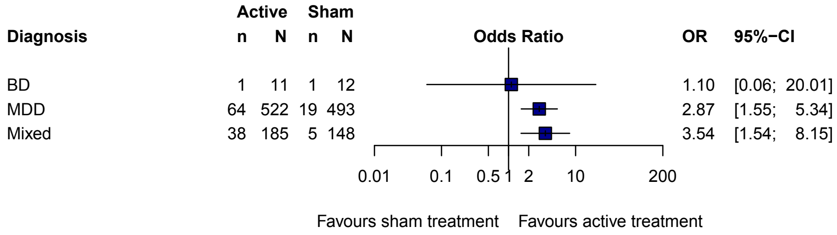


Supplementary Figure 6b. Forest plot of HF-L (exclusion psychosis).

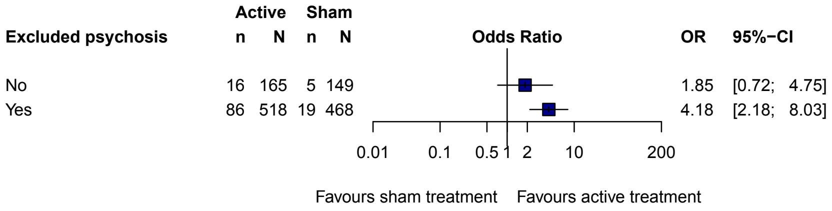


Supplementary Figure 6c. Forest plot of HF-L (hospitalisation status).

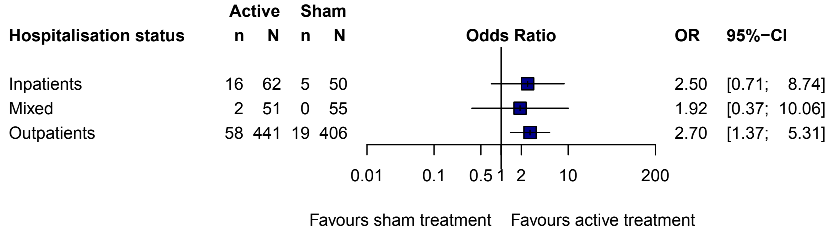


Supplementary Figure 6d. Forest plot of HF-L (treatment resistance).

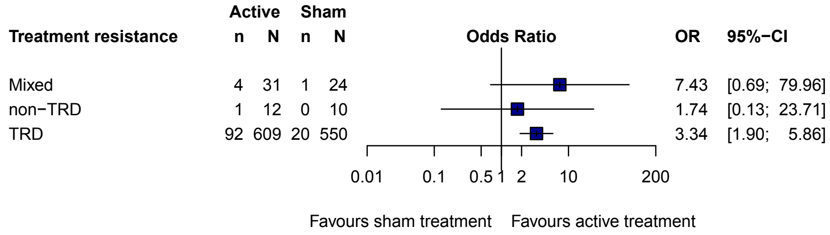

Supplementary Figure 7a. Forest plot of BL (diagnosis).

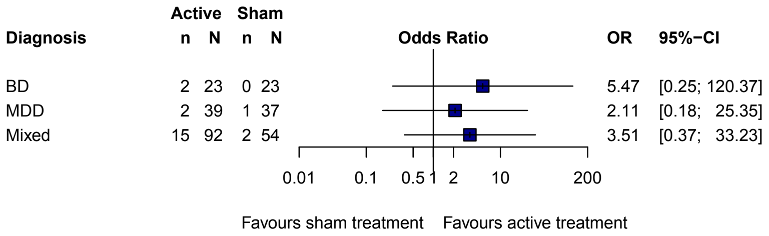


Supplementary Figure 7b. Forest plot of BL (exclusion psychosis).

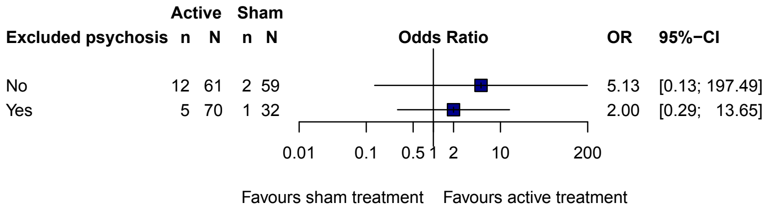


Supplementary Figure 8. Forest plot of tDCS (treatment resistance).

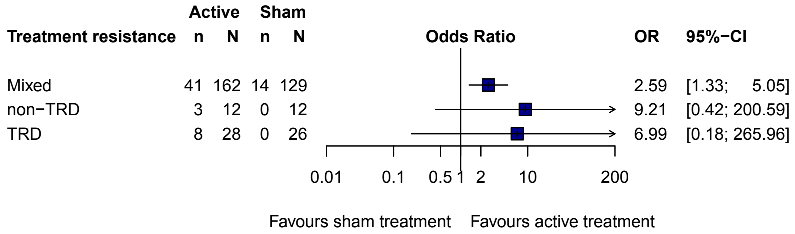

#### 6. Reasons for excluding full-texts

| Did not meet age criteria |
|---|
| Beynel et al., 2014 |
| Blumberger et al., 2012 |
| Chistyakov et al., 2015 |
| Dolberg et al., 2002 |
| Garcia-Toro et al., 2006 |
| He et al., 2011 |
| Höppner et al., 2003 |
| Kang et al., 2016 |
| Kauffmann et al., 2004 |
| Klein et al., 1999 |
| Koerselman et al., 2004 |
| Loo et al., 2003 |
| Loo et al., 2007 |
| Loo et al., 2012 |
| Loo et al., 2017 |
| Manes et al., 2001 |
| Miniussi et al., 2005 |
| Mogg et al., 2008 |
| Mosimann et al., 2004 |
| Nadeau et al., 2014 |
| Padberg et al., 2002 |
| Palm et al., 2012 |
| Plewnia et al., 2014 |
| Rossini et al., 2005 |
| Stern et al., 2007 |
| Triggs et al., 2010 |
| **Different stimulation technique** |
| Barclay & Barclay, 2014 |

| |
|---|
| Carpenter et al., 2017 |
| Fang et al., 2016 |
| Martiny et al., 2010 |
| McClure et al., 2015 |
| Rong et al., 2012 |
| Schutter et al., 2009 |
| Shiozawa et al., 2015 |
| **Did not present data on depressive symptoms** |
| Aguirre et al., 2011 |
| Boggio et al., 2007 |
| Grisaru et al., 1998 |
| Kozel et al., 2011 |
| Minichino et al., 2014 |
| Möller et al., 2006 |
| Nejati et al., 2017 |
| Pascual-Leone et al., 1996 |
| Praharaj et al., 2009 |
| Schutter & Koerselman, 2012 |
| Speer et al., 2009 |
| Speer et al., 2001 |
| Szuba et al., 2001 |
| **Presented duplicate data** |
| Baeken et al., 2015 |
| Baeken et al., 2014 |
| Dang et al., 2007 |
| Hausmann et al., 2004 |
| Herbsman et al., 2009 |
| Lisanby et al., 2009 |
| Loo et al., 2001 |
| Nahas et al., 2001 |
| Powell et al., 2014 |

| |
|---|
| Rosenquist et al., 2013 |
| Schutter et al., 2010 |
| Solvason et al., 2014 |
| Ullrich et al., 2013 |
| **Co-initiation of medication** |
| Bennabi et al., 2015 |
| Hausmann et al., 2004 |
| Herwig et al., 2007 |
| Herwig et al., 2003 |
| Hoeppner et al., 2010 |
| Peng et al., 2012 |
| Ray et al., 2011 |
| Ullrich et al., 2012 |
| Zheng et al., 2015 |
| **Co-initiation of CCT** |
| Brunoni et al., 2014 |
| Segrave et al., 2014 |
| Vanderhasselt et al., 2015 |
| **Co-initiation of sleep deprivation** |
| Krstic et al., 2014 |
| **Did not include a sham condition**[*] |
| Arns et al., 2010 |
| Chistyakov et al., 2010 |
| Fujita & Koga, 2005 |
| Janicak et al., 2010 |
| Kolbinger et al., 1995 |
| Kuroda et al., 2006 |
| Levkovitz et al., 2009 |
| Nongpiur et al., 2011 |
| Rybak et al., 2005 |
| Schrijvers et al., 2012 |

| |
|---|
| Tamas et al., 2007 |
| Vanderhasselt et al., 2009 |
| Vanderhasselt et al., 2016 |
| Woźniak-Kwaśniewska et al., 2015 |
| **Case report** |
| Cohen et al., 2008 |
| Vedeniapin et al., 2010 |
| **Editorial** |
| Lisanby, 2003 |
| **Study protocol** |
| Pereira Junior et al., 2015 |
| **Depression not primary diagnosis** |
| Carretero et al., 2009 |

*Note.* Full-text articles excluded. *for cross-over trials that included a sham condition, data were not available separately for the active and sham conditions prior to the cross-over.

### 7. Risk of bias assessment

*Supplementary Table 2. Cochrane risk of bias tool.*

| | Random sequence generation | Allocation concealment | Blinding of participants and personnel | Blinding of outcome assessment | Incomplete outcome | Selective outcome reporting | Overall risk |
|---|---|---|---|---|---|---|---|
| tDCS | | | | | | | |
| Fregni et al. 2006a | Unclear | Unclear | Unclear | Low | Unclear | Unclear | Unclear |
| Fregni et al. 2006b | Unclear | Unclear | Unclear | Low | Unclear | Unclear | Unclear |
| Boggio et al. 2008 | Unclear | Unclear | Unclear | Low | Low | Low | Unclear |
| Loo et al. 2010 | Unclear | Unclear | Low | Low | Low | Low | Unclear |
| Blumberger et al. 2012 | Low | Low | Unclear | Low | Low | Low | Unclear |
| Brunoni et al. 2013 | Low | Low | Low | Unclear | Low | Low | Unclear |
| Salehinejad et al. 2015 | Unclear | Unclear | Unclear | Unclear | Low | Low | Unclear |
| Salehinejad et al. 2017 | Unclear | Unclear | Unclear | Unclear | Unclear | Low | Unclear |
| Brunoni et al. 2017 | Low | Unclear | Low | Low | Low | Low | Low |
| TMS | | | | | | | |
| Anderson et al., 2007 | Unclear | Low | High | Unclear | Low | Low | High |
| Avery et al., 1999 | Unclear | Unclear | Unclear | Unclear | Low | Low | Unclear |
| Avery et al., 2006 | Low | Unclear | Low | Unclear | Low | Low | Unclear |
| Baeken et al., 2013 | Low | Unclear | Unclear | Low | Low | Low | Unclear |
| Bakim et al., 2013 | Low | Unclear | Unclear | Low | Low | Low | Unclear |
| Berman et al., 2000 | Unclear | Unclear | Low | Low | Low | Low | Unclear |
| Beynel et al., 2014 | Low | Unclear | Low | Low | Low | Low | Low |
| Bortolomasi et al., 2007 | Unclear | Unclear | Unclear | Low | Low | Low | Unclear |
| Boutros et al., 2002 | Low | Unclear | High | Low | Low | Low | High |
| Chen et al., 2013 | Unclear | Unclear | Unclear | Unclear | Low | Low | Unclear |
| Chistyakov et al., 2015 | Unclear | Unclear | Low | Low | Low | Low | Unclear |
| Concerto et al., 2015 | Unclear | Unclear | Unclear | Unclear | Low | Low | Unclear |
| Duprat et al., 2016 | Low | Unclear | Unclear | Unclear | Low | Low | Unclear |
| Eschweilier et al., 2000 | Unclear | Unclear | Unclear | Low | Low | Low | Unclear |
| Fitzgerald et al., 2003 | Unclear | Low | Low | Low | Low | Low | Unclear |
| Fitzgerald et al., 2006 | Low | Low | Low | Low | Low | Low | Low |
| Fitzgerald et al., 2012 | Unclear | Unclear | Unclear | Low | Low | Low | Unclear |
| Garcia- Toro et al., 2001 | Unclear | Unclear | Unclear | Low | Low | Low | Unclear |
| George et al., 1997 | Unclear | Unclear | Unclear | Low | Low | Unclear | Unclear |

| | | | | | | | |
|---|---|---|---|---|---|---|---|
| George et al., 2000 | Unclear | Unclear | Low | Low | Low | Low | Unclear |
| George et al., 2010 | Low | Unclear | Low | Low | Low | Low | Low |
| Hansen et al., 2004 | Low | Unclear | Low | Low | High | Low | High |
| Hernandez- Ribas et al., 2013 | Unclear | Unclear | Unclear | Low | Low | Low | Unclear |
| Holtzheimer et al., 2004 | Unclear | Unclear | Unclear | Low | Low | Low | Unclear |
| Jakob et al., 2008 | Unclear | Unclear | Unclear | Unclear | Low | Low | Unclear |
| Januel et al., 2006 | Unclear | Unclear | Unclear | Low | Low | Low | Unclear |
| Jin and Phillips, 2014 | Low | Unclear | Low | Unclear | Low | Low | Unclear |
| Kimbrell et al., 1999 | Unclear | Unclear | Unclear | Low | Low | Low | Unclear |
| Kreuzer et al., 2015 | Low | Unclear | Low | Low | High | Low | High |
| Leuchter et al., 2015 | Low | Unclear | Low | Low | Low | Low | Low |
| Levokovitz et al., 2015 | Low | Low | Low | Low | Low | Low | Low |
| Li et al., 2014 | Unclear | Unclear | Unclear | Low | Low | Low | Unclear |
| Lingeswaran et al., 2011 | Low | Low | Low | Low | Unclear | Low | Unclear |
| Loo et al., 1999 | Unclear | Unclear | Low | Low | Low | Low | Unclear |
| Loo et al., 2007 | Low | Unclear | Low | Low | Low | Low | Low |
| McDonald et al., 2006 | Unclear | Unclear | Unclear | Low | Low | Low | Unclear |
| Mogg etal., 2008 | Low | Low | High | High | Low | Low | High |
| Nahas et al., 2003 | Low | Unclear | Low | Low | Low | Low | Low |
| O'Reardon et al., 2007 | Unclear | Unclear | Unclear | Low | Low | Low | Unclear |
| Padberg et al., 1999 | Unclear | Unclear | Unclear | Low | Low | Low | Unclear |
| Paillere-Martinot et al., 2010 | Low | Low | Low | Low | Low | Low | Low |
| Pallanti et al., 2010 | Low | Low | Unclear | Low | Low | Low | Unclear |
| Prasser et al., 2015 | Unclear | Unclear | Unclear | Low | Low | Low | Unclear |
| Speer et al., 2014 | Unclear | Unclear | Unclear | Low | Low | Low | Unclear |
| Su et al., 2005 | Unclear | Unclear | Unclear | Low | Low | Low | Unclear |
| Tavares et al., 2017 | Low | Low | Low | Low | Low | Low | Low |
| Theleritis et al., 2017 | Low | Low | Low | Low | Low | Low | Low |
| Zheng et al., 2010 | Unclear | Unclear | Unclear | Unclear | Low | Low | Unclear |
